# Changes in Microorganisms in the Rhizosphere of Mulberry Genotypes with Differing Resistance to Bacterial Wilt

**DOI:** 10.1101/407494

**Authors:** Yao Guo, Cui Yu, Xingming Hu, Wen Deng, Yong Li, Zhixian Zhu, Rongli Mo, Zhaoxia Dong

## Abstract

A close relationship between soil-borne diseases, soil microbial community structure, and functional diversity has been described in the mulberry plant. In the present study, microbial abundance, community structure, and functional diversity in the soil rhizosphere were compared in resistant (Kangqing10) and susceptible (Guisang12) mulberry genotypes using the dilution plate method, micro-ecology technology, and polymerase chain reaction-denaturing gradient gel electrophoresis (PCR-DGGE). The goal of this study was to develop better management methods for mulberry cultivation and preventing and controlling bacterial wilt. Rhizosphere soil microorganisms were more abundant in the resistant normal mulberry genotype than in the susceptible normal mulberry genotype. Carbon source utilization was better in the susceptible normal mulberry genotype. These properties were lower in the resistant sickly mulberry genotype than in the susceptible sickly mulberry genotype. PCR-DGGE indicated that the bacterial and fungal community structures of the resistant genotypes were more stable than those of the susceptible genotypes. Correlation regression analysis implicated mulberry bacterial wilt in the loss of soil nutrients, particularly organic matter and nitrogen, which can disrupt the balance of the soil microbial community. Loss of soil organic matter and nitrogen had a lower impact on resistant genotype plants than on susceptible genotype plants. Therefore, resistant genotype plants displayed some resistance to bacterial wilt. Further insights into the soil rhizosphere microbial diversities of resistant and susceptible genotypes will help in the control and prevention of mulberry bacterial wilt.

## Introduction

Bacterial wilt is a devastating soil-borne disease caused by Ralstonia solanacearum and intestinal bacteria. It is widely distributed in tropical, subtropical, and temperate regions [1] and is common in commercial crops including Solanaceae [2], Leguminosae [3], tobacco [4], Cucurbitaceae [5], and Sanko [6-8]. One of the areas that has been affected is Guangxi, a key silkworm area in China. Guangxi accounts for more than 30% of the national mulberry area. However, the extension of mulberry planting years and improper cultivation management have led to a continuous annual decrease in the yield of mulberry leaves. Feeding of silkworms requires the collection of mulberry leaves every year to ensure that the mulberry supply exceeds demand. Thus, farmers tend to apply inorganic fertilizer, particularly nitrogenous fertilizer, over long periods to increase yield. Although this practice can increase production, in the long-term it will alter the physical and chemical properties of the soil [9], reduce soil microbial diversity [10], and promote the development of severe soil-borne diseases such as bacterial wilt [11, 12]. As a result, the stability and sustainable development of the silk industry may be seriously threatened by the considerable loss of mulberry plants or yield of mulberry leaves.

Several recent studies have suggested that primary self-regulation of the soil micro-ecological environment by adding organic fertilizer can help prevent soil-borne diseases. Application of organic fertilizer can improve soil microbial community composition, enrich soil microbial community structures, and increase the quantities of root exudates, thus significantly reducing the incidence of bacterial wilt [13-18]. Organic fertilizer is applied to prevent and control soil-borne diseases of various vegetables [16] and strawberries in greenhouse environments [13]. The occurrence of tomato bacterial wilt can be controlled and tomato yield increased by adding silicon fertilizer and bagasse [18].

Another approach involves the use of a balanced soil bacterial community structure using different cropping systems tocontrol bacterial wilt. The incidence of tobacco bacterial wilt can be significantly reduced by rotating tobacco with corn, lily, and carrot [19]. A third approach involves biological control to reduce the rate of bacterial wilt. For example, an endophytic Bacillus subtilis strain can prevent mulberry bacterial wilt [20]. Despite these approaches, further improvements have been limited by gaps in knowledge. The microbial community structure in the rhizosphere surrounding the plant roots, as well as the changes in the microbial community that promote bacterial wilt, is not well-understood. To explore these issues, we carried out a study in Guangxi province, the site of the largest sericulture production in China. Our results revealed the diversity of the rhizosphere microbial community structure between resistant and susceptible genotypes and their association with bacterial wilt, increasing the understanding of the micro-ecological mechanisms of bacterial wilt in the surrounding rhizosphere and conditions that improve resistance to bacterial wilt. These results will help improve the prevention and control of mulberry bacterial wilt.

## Material and Methods

### Collection and preparation of soil samples

Eight soil samples were collected froma mulberry breeding site in Xinsheng Village, Gula town, Binyang County, Guangxi province (23°06’N,108°59’E) in September 2017. All mulberry trees were 7Y cultivations. The rhizosphere soil of the resistant and susceptible mulberry genotypes was both sandy loam. Urea-based fertilizer was applied twice per year (1100 kg/hm2 each time), with 40% applied in the spring (March) and 60% after the summer harvest in June. Soil rhizosphere samples were collected using the five-point sampling method [21] and shaking root method [22]. The control group, which was designated as CK, comprised rhizosphere soilfrom the base ofnormal (including resistant and susceptible genotypes) mulberry plants. Rhizosphere soil was also acquired from the base of sickly (including resistant and susceptible genotypes) mulberry plant genotypes (T1, T2, and T3). Additional information is presented in Table 1. Each obtained rhizosphere soil was placed in three labeled and sealed bags. One sample was partially dried for physical and chemical analyses. The other two samples were stored at 4°C or −20°C for microbial analysis.

**Table 1:** Designation of soil samples. Resistant normal mulberry genotype is denotedas RN (resistance normality), susceptible normal mulberry genotype is denotedas SN (susceptibility normality), resistant sickly mulberry genotypeis denoted as RS (resistance sickness), and susceptible sickly mulberry genotypeis denotedas SS (susceptibility sickness).

| Soil sample numbers and variety names |  |  |
| --- | --- | --- |
| Normality | CK | RN (Kangqing10, normal and control group) |
|  |  | SN (Guisang12, normal and control group) |
| Sickness | T1 | RS1 (Kangqing10-1 group) |
|  |  | SS1 (Guisang 12-1 group) |
|  | T2 | RS2 (Kangqing 10-2 group) |
|  |  | SS2 (Guisang 12-2 group) |
|  | T3 | RS3 (Kangqing 10-3 group) |
|  |  | SS3 (Guisang 12-3 group) |

### Physicochemical properties of rhizosphere soil

The physicochemical properties of the rhizosphere soil, including alkaline nitrogen, available phosphorus, available potassium, and soil organic matter, can reflect the basic fertility status of the tested soil. The physicochemical properties of the collected rhizosphere soil are shown in Table 2. Alkaline N was extracted with 1.07M NaOH by the alkali-hydrolysis and diffusion method [23]. Available P was extracted with 0.5 M NaHCO_3_ as previously described [23, 24]. Available K was extracted with 1M NH_4_OAc (1:10 soil:solution ratio) for 1h and analyzed by atomic absorption spectrophotometry [23, 25]. Soil organic matter was extracted with 0.8M K_2_Cr_6_O_7_ using the standard Walkley-Black potassium dichromate oxidation method [23, 26].

**Table 2:** Comparison of physicochemical properties in soil rhizosphere with different resistant mulberry genotypes. The genotypes included Kangqing10, normal and control group (RN), Guisang12, normal and control group (SN), Kangqing10-1 group (RS1), Guisang 12-1 group (SS1), Kangqing 10-2 group (RS2), Guisang 12-2 group (SS2), Kangqing 10-3 group (RS3), and Guisang 12-3group (SS3). †

|  |  | Alkaline N<br>(mg kg<sup>-1</sup>) | Available P<br>(mg kg<sup>-1</sup>) | Available K<br>(mg kg<sup>-1</sup>) | SOM<sup>‡</sup>(%) |
| --- | --- | --- | --- | --- | --- |
| CK | RN | 82.25±0.35<sup>a</sup> | 44.59±15.09<sup>b</sup> | 35.00±0.00<sup>e</sup> | 4.26±0.05<sup>a</sup> |
|  | SN | 53.55±0.35<sup>d</sup> | 96.36±19.94<sup>a</sup> | 102.50±2.50<sup>b</sup> | 2.69±0.00<sup>d</sup> |
| T1 | RS1 | 71.75±3.15<sup>b</sup> | 49.77±3.97<sup>b</sup> | 52.50±2.50<sup>c</sup> | 3.83±0.05<sup>b</sup> |
|  | SS1 | 65.80±0.70<sup>c</sup> | 27.52±0.44<sup>c</sup> | 40.00±2.50<sup>d</sup> | 3.23±0.05<sup>c</sup> |
| T2 | RS2 | 11.55±1.05<sup>g</sup> | 1.41±0.11<sup>e</sup> | 10.00±0.00<sup>g</sup> | 0.48±0.00<sup>g</sup> |
|  | SS2 | 8.40±0.70<sup>h</sup> | 5.93±1.54<sup>e</sup> | 41.25±1.25<sup>d</sup> | 0.45±0.05<sup>g</sup> |
| T3 | RS3 | 30.80±0.00<sup>e</sup> | 21.57±1.54<sup>cd</sup> | 122.50±2.50<sup>a</sup> | 0.69±0.03<sup>f</sup> |
|  | SS3 | 26.60±0.70<sup>f</sup> | 7.91±0.88<sup>de</sup> | 15.00±0.00<sup>f</sup> | 0.79±0.01<sup>e</sup> |

<sup>†</sup>Data are expressed as the mean ± SD (n = 3). The superscript letters that differ within a column indicate significant differences between treatments (P < 0.05).
<sup>‡</sup>Soil organic matter (SOM).

### Viability of cultured bacteria and actinomycetes

The total numbers of viable cultured bacteria and actinomycetes were both measured as colony forming units (CFUs) on agar plates using the dilution plate method [27]. The media used to separately culture bacteria and actinomycetes werebeef extract peptone medium and Gause’s No.1 synthetic medium, respectively [28]. No antibiotics were present. The sample dilution used for culture was 5g of each fresh soil sample in 45mL sterilized 0.85% NaCl. Each dilution was shaken for 15 min at 150 rpm and then left standing for 30 min. The individual bacterial and actinomycete strains from the 1×10^-4^ and 1×10^-3^ dilution plate, respectively, were isolated and purified. The purified strains were cultured on Biolog recommended agar media (BUG agar; Biolog, Inc., Hayward, CA, USA) and incubated at 25°C in the dark. Pure cultures from the BUG agar were then grown on GEN III MicroPlates (Biolog) for 24-72h and analyzed using a Biolog MicroStation equipped with Microlog microbial identification software (Ver. 5.2.01). The identification results were accepted if the similarity index value was ≥ 0.5. Three replicates were analyzed for each sample.

### Microbial community structure diversity analysis and microbial identification Soil microbial community diversity

Ninety-six-well microplates (Ecoplates; Biolog, Inc.) were used to analyzesoil microbial community diversity based on the utilization of 31 carbon sources as previously described [29]. The culture dilution was obtained as described above. The suspensionwas obtained from 1×10^-2^ dilutions after 30 min. Each suspension was dispensed into thewells of Ecoplates (125mL per well) with a multi-channel repetitive-dispensing pipette. The plates were incubated at 25°C in the dark and absorbance at 590 and 750 nm was recorded every 24h for 7 days using the reader incorporated into the GEN III microStation. Three replicates per treatment and sampling time were analyzed.

Each well of an Ecoplate was loaded with a single carbon source. Thirty-one single carbon sources belonging to six classes (carbohydrates, amino acids, carboxylic acids, amines, polymers, and miscellaneous) were used in the analysis. The average well color development, which reflects the overall utilization of soil microorganisms on a single carbon source, was calculated as described by Garland JL [30]. Average well color development data from 144h were used to calculate the functional diversities using the Shannon index and Shannon evenness [31, 32]. The Shannon index formula was calculated as H= -∑Pi lnPi,Pi by subtracting the control from each substrate’s absorbance and then dividing this value by the total color change recorded for all 31 substrates. Evenness was calculated as E = H/ln(richness), where richness is the number of substrates utilized[31].

### DNA extraction and PCR amplification of 16s rRNA and ITS

Total DNA was extracted using 0.3g of each fresh soil sample. DNA was extracted using an E.Z.N.A.® Soil DNA kit (Omega Bio-Tek, Inc., Norcross, GA, USA) according to the manufacturer’s instructions. The remaining samples were kept at −20°C for subsequent use. Specific gene fragments of bacteria and fungi were specifically amplified using the extracted DNA sample as a template with appropriate primers. Primers GC(CGCCCGCCGCGCCCCGCGCCCGGCCCGCCGCCCCCGCCCC) + 341F (5’-CCTACGGGAGGCAGCAG-3’) and R518 (5’-ATTACCGCGGCTGCTGG-3’) were used to directly amplify the V3 hypervariable regions of 16s Rrna bacterial sequences from the microbial genome DNA [33, 34]. Amplification of the internal transcribed spacer (ITS) regions of fungal sequences from the microbial genome DNA used primers GC (CGCCCGCCGCGCCCCGCGCCCGGCCCGCCGCCCCCGCCCC) + Fung (5’-ATTCCCCGTTACCCGTTG-3’) and NS1 (5’- GTAGTCATATGCTTGTCTC-3’) [35, 36]. The reaction mixtures (30 μL) contained 15 Ml of 2 x Es Taq MasterMis, 1.2 μL of 10μM forward primer, 1.2 μL of 10μ Mreverse primer, 9.6 μL of RNase-free water, and 3 μL of template DNA. PCR conditions of bacteria were 36 cycles of 94°C for 5 min, 94°C for 1min, 64°C for 1min, and 72°C for 1min; and a final elongation at 72°C for 7 min. PCR conditions of the fungus were 35 cycles of 94°C for 5 min, 94°C for 30s, 47°C °C for 45s, and 72°C for 1min; and a final elongation at 72°C for 10 min. PCR products were excised from a 1.0% agarose gel and quantified in a Qubit1 Fluorometer (Invitrogen, Carlsbad, CA, USA) using a Quant-iTTM dsDNA HS assay kit (Invitrogen).

### Denaturing gradient gel electrophoresis (DGGE) of 16s rRNA and ITS fragments

The amplified bacterial and fungal gene fragments (both 75 ng of DNA) obtained using the V3 hypervariable regions of 16s rRNA and ITS regions, respectively, were separated by DGGE using a polyacrylamide gel containing 8% acrylamide. The D-code system (Bio-Rad, Hercules, CA, USA) was used; this system can separate DNA fragments that differ in sequence but have the same length. Gradients of 50-80% and 20-50% were used for bacteria and fungi, respectively. The cross-linking agent was ammonium persulfate (APS) and N,N,N,N,tetramethyl-ethylene diamine (TEMED) (gradient gel:APS:TEMED = 1000:10:1). Electrophoresis was conducted in 1xTAE buffer at 60C at a constant voltage of 80 V for 18 h. The DGGE gels were stained with SYBR1 Green (Molecular Probes,Eugene, OR, USA) and analyzed using the Quantity One v4.6.2 gel analysis software (Bio-Rad). Bands with intensities <0.05 were excluded from analysis. Two replicates were analyzed for each sample. The Shannon index and Shannon evenness were both calculated as described for the Ecoplate analysis.

### Statistical analyses

The data were mainly analyzed using one-way analysis of variance (Levene’s test was used to assess the equality of variances before performing analysis of variance), and significant differences between the means were determined by the least significant difference test. The differences were considered as significant when P < 0.05. DPS v7.05 software was used to analyze the diversity index and evenness of Ecoplates and DGGE, the differences in the main six types of carbon sources utilization, and their correlations. The absolute value of the correlation coefficient (R) revealed no correlation at R=0-0.09, weak correlation at R=0.1-0.3, moderate correlation at R=0.3-0.5, and strong correlation at R=0.5-1.0. Quantity One v4.6.2 software (Bio-Rad) was used to quantitatively analyze DGGE bands. Excel 2016 software (Microsoft, Redmond, WA, USA) was used to analyze and map the data. The NCBI website was used for BLAST nucleic acid alignment for partial post-recovery sequencing results.

## Results

### Viability of cultured bacteria and actinomycetesin soil rhizosphere of mulberry with different resistant genotypes to bacterial wilt

The abundance of cultured microorganisms differed in the soil rhizosphere of mulberry genotypesthat were resistant and susceptible to bacterial wilt (Fig 1). The numbers of viable bacteria were greater than those of viable actinomycetes in resistant mulberry genotypes (RN, RS1, RS2, RS3) and susceptible mulberry genotypes (SN,SS2, SS2, SS3). The numbers of cultured rhizosphere bacteria and actinomycetes of the resistant normal mulberry genotypes (RN) wereboth significantly higher (P<0.05) than those of the susceptible normal mulberry genotypes (SN). The number of rhizosphere actinomycetes of the resistant sickly mulberry genotypes (RS1,RS2,RS3) was also higher than that of the susceptible sickly mulberry genotypes (SS1,SS2,SS3), except for T1. However, the number in the rhizosphere of the resistant sickly mulberry genotypes (RS1,RS2,RS3) was significantly lower (P<0.05) than those of the susceptible sickly mulberry genotypes (SS1,SS2,SS3), except for T3 bacteria. The numbers of cultured rhizosphere bacteria and actinomycetes of the resistant normal mulberry genotypes (RN) were all significantly higher (P<0.05) than those of the resistant sickly mulberry genotypes (RS1,RS2,RS3), but the number of cultured rhizosphere bacteria and actinomycetes of the susceptible normal mulberry genotypes (SN) were significantly higher (P<0.05) than those of the susceptible sickly mulberry genotypes (SS1,SS2,SS3), except for T3 bacteria. These results indicate that the abundance of soil microorganisms of the RN genotypes in the rhizosphere was higher than that of the SN genotypes, while the number of soil microorganisms of the RS genotypes in the rhizosphere was lower than that of the SS genotypes.

**Fig 1.**
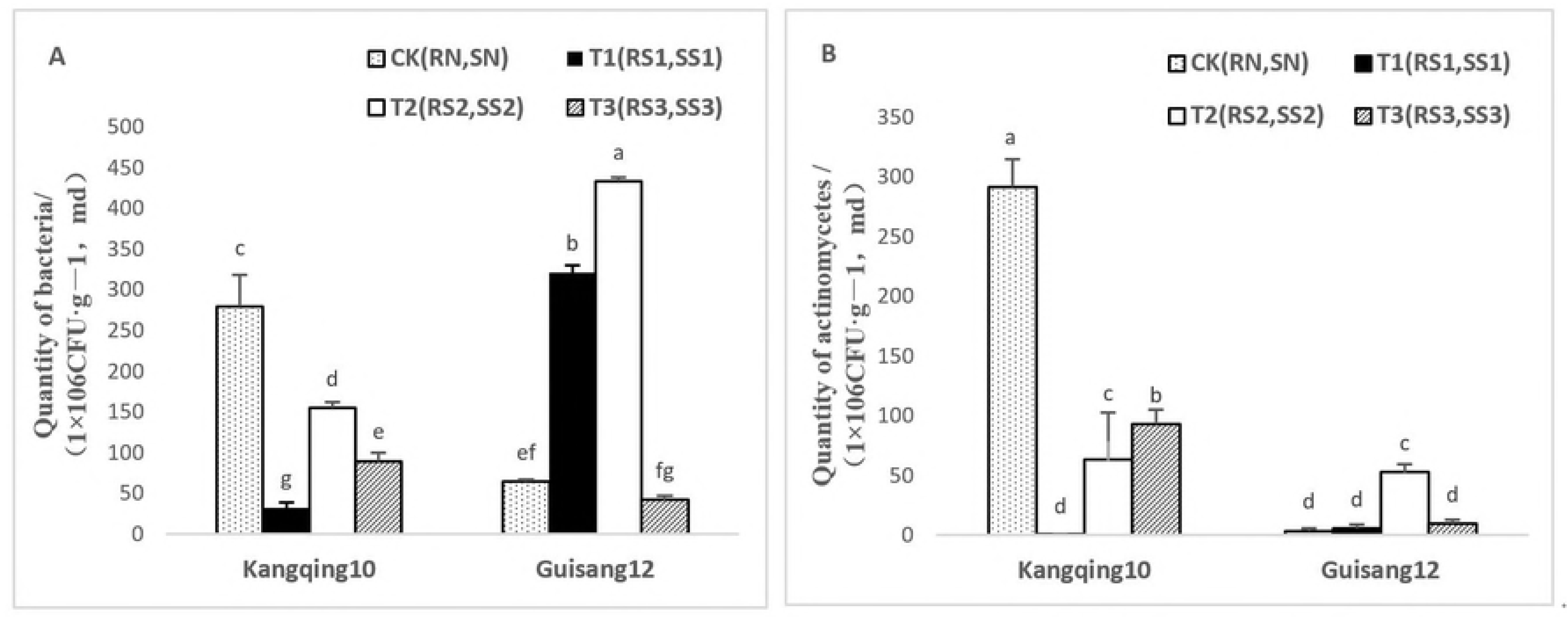
Comparison ofabundance of (A) bacteria and (B) actinomycetes in the soil rhizosphere with different resistant mulberry genotypes treated with CK: Kangqing10, normal and control group (RN), Guisang12, normal and control group (SN); T1: Kangqing10-1 group (RS1), Guisang 12-1 group (SS1); T2: Kangqing 10-2 group (RS2), Guisang 12-2 group (SS2); T3: Kangqing 10-3 group (RS3), Guisang 12-3 group (SS3). The bars indicate the mean ± SD (n = 3). Different letters indicate significant differences between treatments (P < 0.05).

### Average well color development in soil rhizosphere of mulberry with different resistant genotypes to bacterial wilt

The average well color development can reflect the overall utilization of soil microorganisms on a single carbon source. Microbial activity in the soil samples was examined as previously described [29]. During 168h incubation of the soil, the average well color development values of all soil samples increased rapidly starting at 24h and reached a steady state at approximately 144h. In this experiment, 144h was selected for statistical analysis (Fig 2). Soil microbes in the different resistant mulberry genotypes displayed differing utilization of the single carbon source. The equilibrium values of the steady state of single carbon source utilization were larger in susceptible genotypes than in resistant genotypes, suggesting that the rate of carbon source utilization of resistant mulberry genotypes was higher than that of susceptible mulberry genotypes. For soil microbes of resistant mulberry genotypes, RN and RS2 sickly plants showed the highest utilization of the carbon source, followed by RS3 sickly plants, with RS1 sickly plants displaying the weakest utilization of the single carbon source. For soil microbes of the susceptible mulberry genotypes, the SS1 sickly plants displayed the greatest utilization of the single carbon source, followed by SS3 sickly plants, with SN normal plants displaying the weakest utilization of the single carbon source. Additionally, the utilization of carbon sources by RN in the rhizosphere was higher than that of SN. In contrast, utilization of the carbon sources by RS in the rhizosphere was lower than that of susceptible sickly mulberry genotypes in the order of RS1< SS1; RS2< SS2; and RS3<SS3. These results suggest that the microbial function in the resistant normal genotypes in the surrounding rhizosphere was much greater than that of the susceptible normal genotypes, while the microbial function in the resistant sickly rhizosphere genotypes was lower than in the susceptible sickly genotypes.

**Fig 2.**
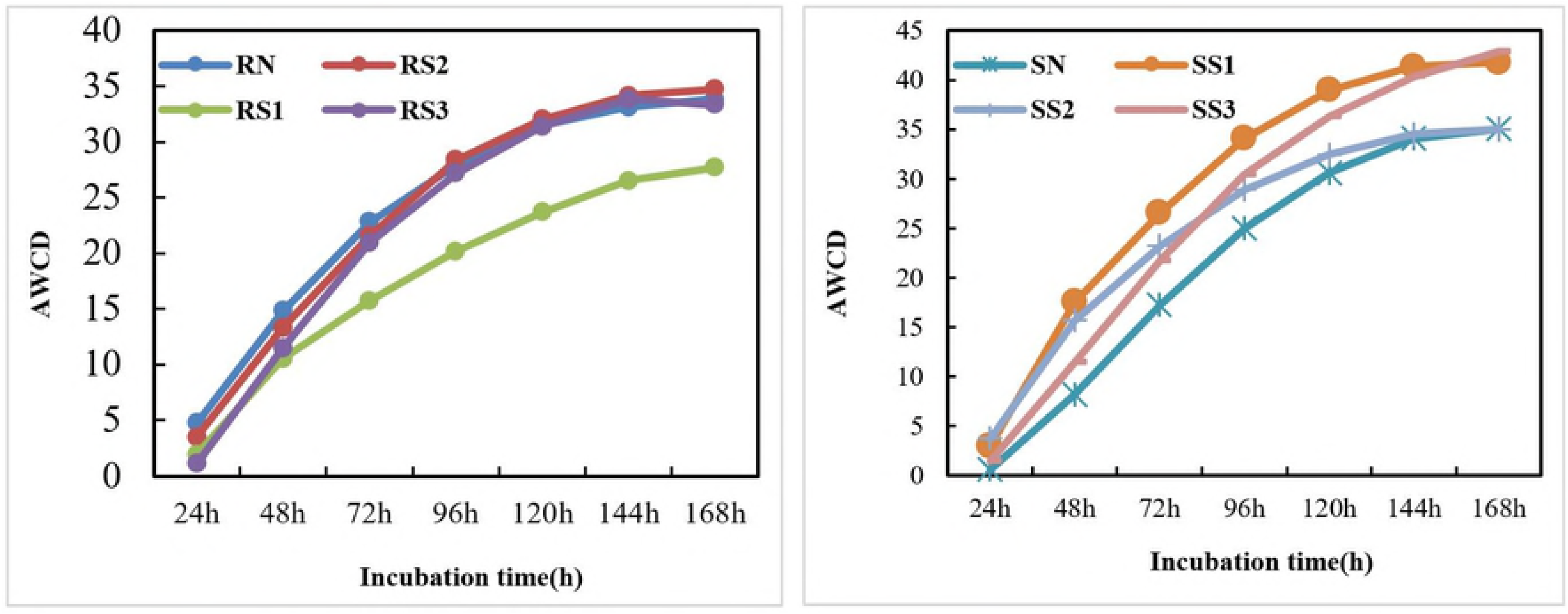
Changes in the average well color development of 31 carbon sources in the soil rhizosphere with different resistant mulberry genotypes treated with Kangqing10, normal and control group (RN); Guisang12, normal and control group (SN); Kangqing10-1 group (RS1); Guisang 12-1 group (SS1); Kangqing 10-2 group (RS2); Guisang 12-2 group (SS2); Kangqing 10-3 group (RS3);and Guisang 12-3 group (SS3).

The Shannon function index reflecting the rhizosphere soil microbe carbon source utilization by RN was greater than that of SN, while the Shannon function index of resistant sickly mulberry rhizosphere genotypes (RS1, RS2, RS3) was less than that of susceptible sickly mulberry genotypes (SS1, SS2, SS3) (Fig 3). The results may reflect the larger numbers of RN genotypes of cultured bacteria and actinomycetes than those of SN genotypes, while the numbers of RS1, RS2, and RS3 rhizosphere bacteria genotypes (RS1, RS2, RS3) were lower than those of the SS1, SS2, and SS3 genotypes, which differed based on the cultivation and fertilization systems in the mulberry garden.

**Fig 3.**
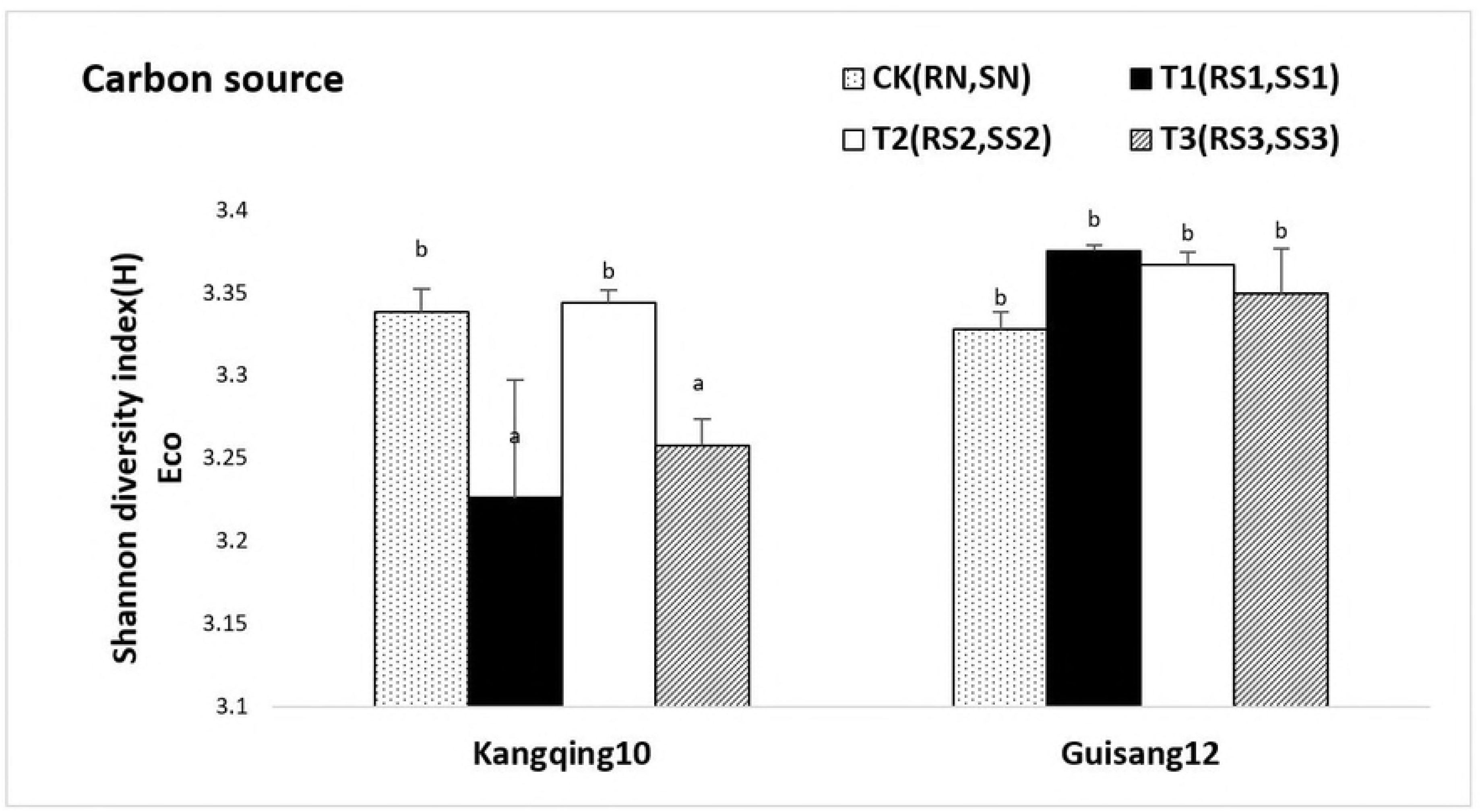
Comparison of the Shannon index (H) of carbon sourcein soilrhizospherewith different resistant mulberry genotypes treated with CK: Kangqing10, normal and control group (RN), Guisang12, normal and control group (SN); T1: Kangqing10-1 group (RS1), Guisang 12-1 group (SS1); T2: Kangqing 10-2 group (RS2), Guisang 12-2 group (SS2); T3: Kangqing 10-3 group (RS3), Guisang 12-3 group (SS3). The bars indicate the mean ± SD (n = 3). Different letters indicate significant differences between treatments (P < 0.05).

In general, RN genotypes displayed the greatest utilization of the single carbon source, followed by SS genotypes, with SN genotypes most weakly utilizing the single carbon source. The data indicate that the utilization of a single carbon source in rhizosphere soil was greater inresistant mulberry genotypes and lower in susceptible mulberry genotypes.

### Classes of carbon sources in soil rhizosphere of mulberry with different resistant genotypes to bacterial wilt

The six classes of carbon sources in soil rhizosphere showed a sequence of utilization for resistantand susceptible mulberry genotypes. The order of utilization was carbohydrates > carboxylic acids > amino acids > polymers > amines > miscellaneous (Fig 4). Amino acids, amines, polymers, and miscellaneous utilization of the RN rhizosphere genotypes were all higher than that of the SN genotypes, while utilization of carbohydrates and carboxylic acidswas lower than that of the SN genotypes. Utilization of carbohydrates, carboxylic acids, amino acids, and miscellaneous molecules by the RS rhizosphere genotypes was all lower than those of the SS genotypes, while utilization of amines and polymers utilization was higher than that of the SS genotypes. Furthermore, utilization of carboxylic acids, amino acids,polymers, andmiscellaneous molecules by the RN rhizosphere genotypes was higher than that of the RS1, RS2, and RS3 genotypes, with the exceptions of T2 amino acids, polymers, and carboxylic acids. Moreover, they were all significantly different from RS1 (P < 0.05), and utilization of carboxylic acids and polymers differedsignificantlyin RS3 (P<0.05). However, the utilization of carbohydrates and aminesof the RN rhizosphere genotypes was lower than that of the RS1, RS2, and RS3 genotypes, except for T1 amines and T2 carbohydrates; both displayed a significant difference from RS3 (P <0.05). In comparison, the utilization of all six classesof carbon sourcesby the SN rhizosphere genotypes was lower than the utilization by the SS1, SS2, and SS3 genotypes, except for T2 amines and carbohydrates; additionally, utilization of the polymers was significantly different from SS1, SS2, and SS3 (P < 0.05). The utilization of carbohydrates, carboxylic acids, and miscellaneousmolecules displayed significant differencesfrom SS1 and SS3(P < 0.05), and utilization.

**Fig 4.**
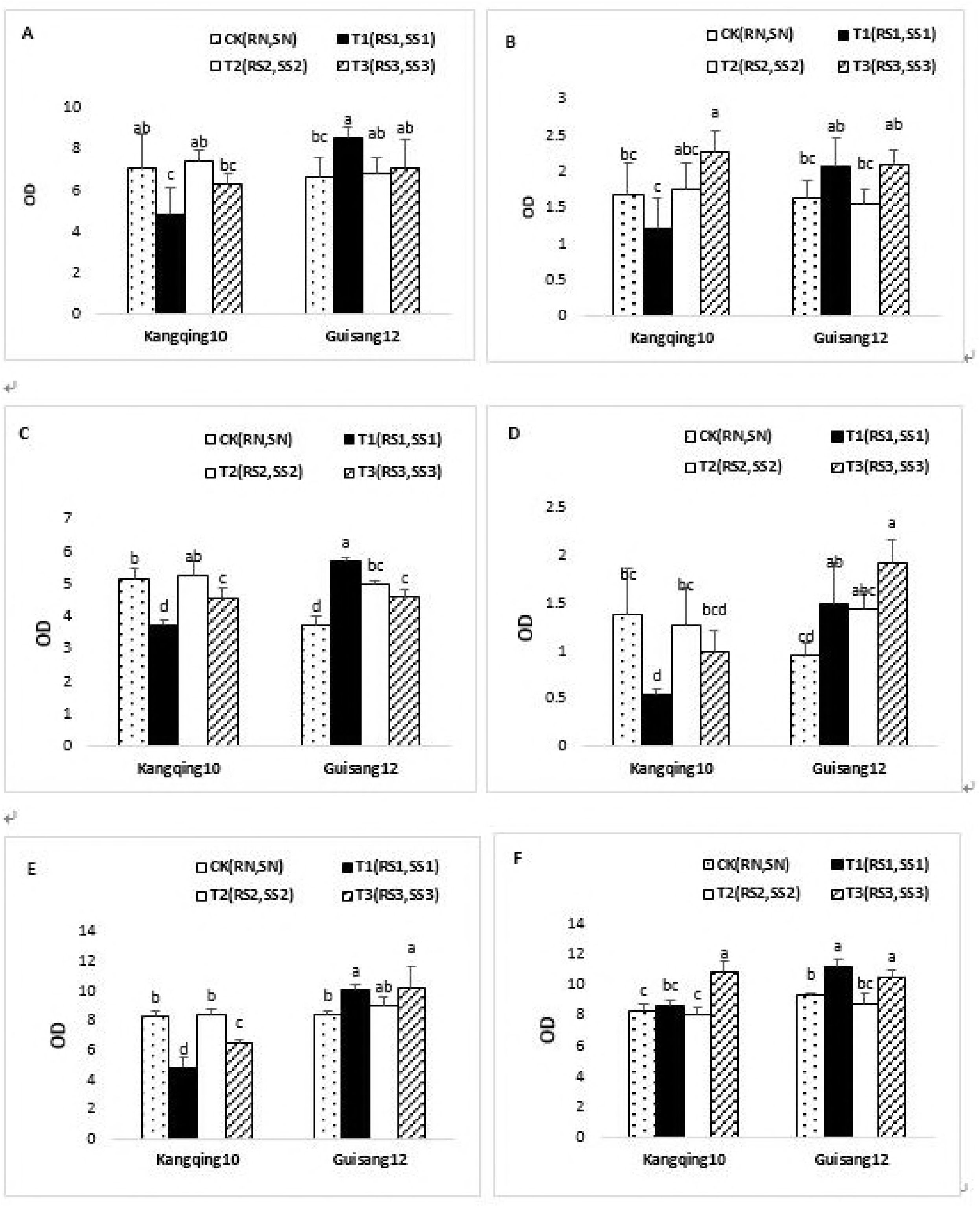
Comparison ofthe six classes of carbon sources in soil rhizosphere with different resistant mulberry genotypes treated with CK: Kangqing10, normal and control group (RN), Guisang12, normal and control group (SN); T1: Kangqing10-1 group (RS1), Guisang 12-1 group (SS1); T2: Kangqing 10-2 group (RS2), Guisang 12-2 group (SS2); T3: Kangqing 10-3 group (RS3), Guisang 12-3 group (SS3). Substrates: (A) amino acids, (B) amines, (C) polymers, (D) miscellaneous, (E) carboxylic acids, and (F) carbohydrates. The bars indicate the mean ± SD (n = 3). Different letters indicate significant differences between treatments (P < 0.05).

### PCR and recycling initial inspection

PCR electrophoresis bands of bacteria and fungi were pronounced, indicating the high accuracy of the DGGE profiles (Fig 5). The recycling electrophoresis bands of bacteria and fungi were also quite pronounced, also indicating the high accuracy of partial sequencing and NCBI alignment.

**Fig 5.**
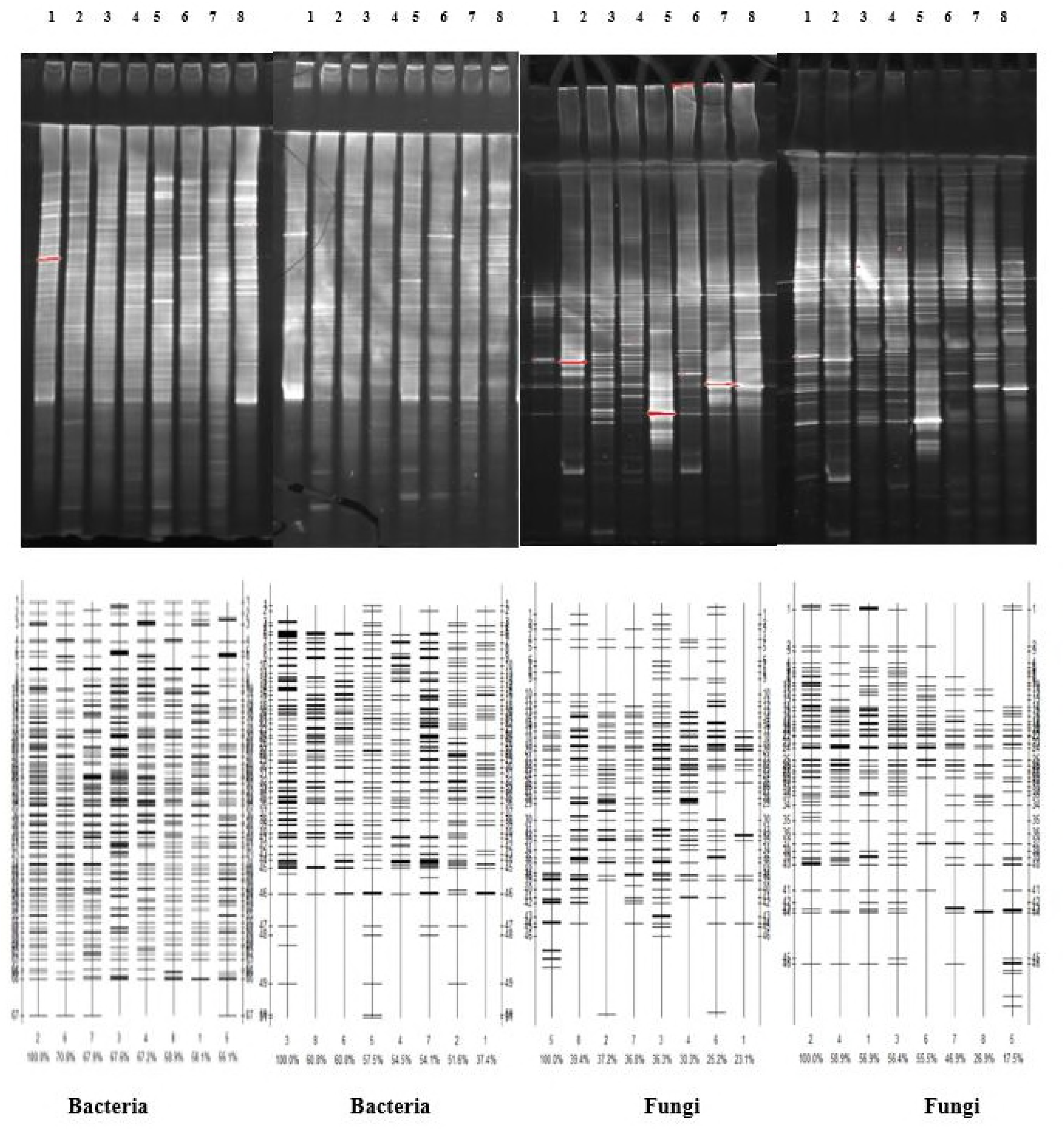
Comparison of the PCR and recycling initial inspection of bacteria and fungi in soil rhizosphere with different resistant mulberry genotypes treated with Kangqing10 normal and control group (RN); Guisang12, normal and control group (SN); Kangqing10-1 group (RS1); Guisang 12-1 group (SS1); Kangqing 10-2 group (RS2); Guisang 12-2 group (SS2); Kangqing 10-3 group (RS3); Guisang 12-3 group (SS3). (A) Bacterial productions of PCR amplification. (B) Bacterial productions of recycling initial inspection. (C) Fungal productions of PCR amplification. (D) Fungal productions of recycling initial inspection.

### DGGE profiles analysis

Differences in DGGE profiles were evident between bacteria and fungi in the eight soil samples (Fig 6). Bands appearing in the bacteria DGGE profiles were significantly differentfrom those in the fungi DGGE profiles; all were higher than thosein the fungi DGGE profiles. The bands of bacteria and fungi of the RN rhizosphere genotypes were all more prominent that those of the RS1, RS2, and RS3 genotypes. The bands for bacteria and fungi of the SN rhizosphere genotypes were more prominent than those for the SS1, SS2, and SS3 genotypes. These results indicate that the microorganism community structures of normal genotype plants were richer than those of the sickly genotype plants. However, there was no obvious regularitybetween the bands of resistant and susceptible genotypes, which may be associated withthe soil nutrients or cropping systems used.

**Fig 6.**
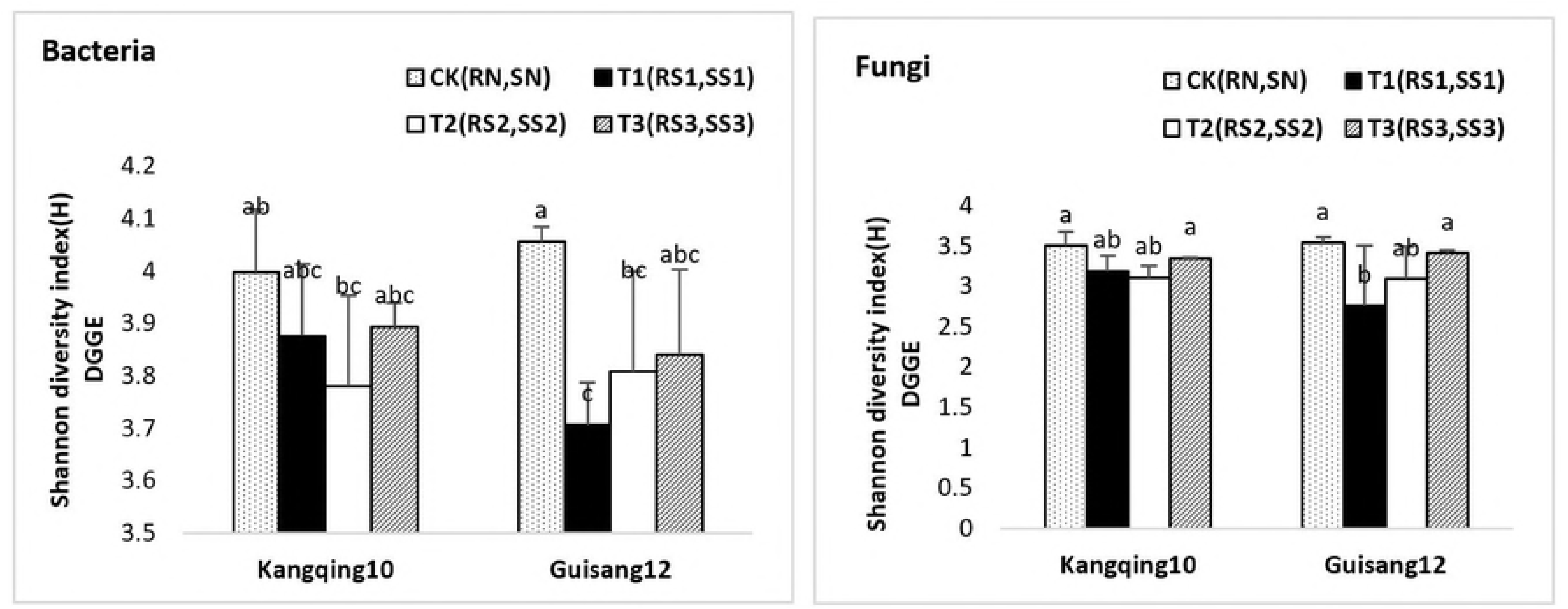
DGGE profiles and results of the analysis of bacteria and fungi in soil rhizosphere with different resistant mulberry genotypes treated with 1-SS1 (Guisang 12-1 group) 2-RN (Kangqing10, normal and control group); 3-SN (Guisang12, normal and control group); 4-SS3 (Guisang 12-3 group); 5-RS3 (Kangqing 10-3 group); 6-RS2 (Kangqing 10-2 group); 7-RS1 (Kangqing10-1 group); 8-SS2 (Guisang 12-2 group).

### Microbial community structure diversity index and evenness analysis in DGGE technology

The Shannon indices of microbial community structure are presented in Fig 7. The Shannon index of bacteria community structure of the RN rhizosphere genotypes waslower than that of the SN genotypes (SN). The Shannon index of bacterial community structure of the RS1, RS2, and RS3 rhizosphere genotypes was larger than those of theSS1, SS2, and SS3 genotypes, which wasopposite to the Shannon index of soil microbial carbon source utilization. The Shannon index of the bacterial community structure of the RN rhizosphere genotypes was larger than those of the RS1, RS2, and RS3 genotypes, which is consistent with the Shannon index of soil microbial carbon source utilization. However, the Shannon index of bacterial community structure of the SN rhizosphere genotypes was larger than that of the (SS1, SS2, SS3 genotypes, with a significant difference inSS1 and SS2 (P<0.05). These results contrasted those of the Shannon index of the soil microbial carbon source utilization. In contrast, the Shannon index of the fungal community structure of the RS1, RS2, and RS3 rhizosphere genotypes was lower than those of the SS1, SS2, and SS3 genotypes, which is consistent with the Shannon index of soil microbial carbon source utilization. The results for the other compared groups (RN with SN, RN with RS, and SN with SS) were also consistent with the Shannon index of bacterial community structure, which was consistent with the Shannon index of soil microbial carbon source utilization. Furthermore, for all Shannon index values of bacterial and fungal community structure, the Shannon index of the SN genotypes was largest, followed by the Shannon index of the RN genotype, with the Shannon index of the SS genotypes showing the lowest value. These results contrast those of the Shannon index of soil microbial carbon source utilization. For susceptible genotypes, although the microbial community structure in normal genotype plantsis richer than that in sickly genotype plants, the ability to utilize carbon sourcesispoor. The microbial community structure, particularly bacteria, is easily altered. In contrast, the microbial community structure in resistant genotypes, which better utilize the carbon, ismore stable than that in susceptible genotypes. Thus, the resistant genotypes exhibit some resistance to bacterial wilt.

**Fig 7.**
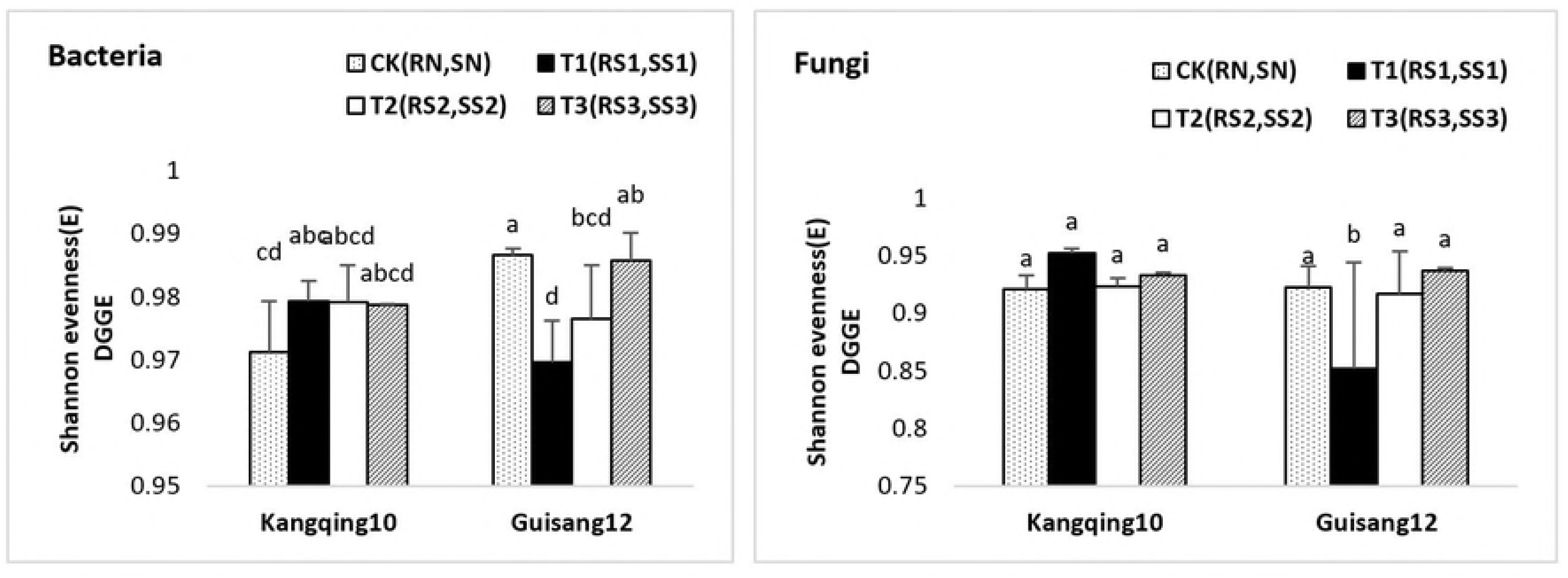
Comparison ofthe Shannon index (H) of bacterial and fungal community structure in soil rhizosphere with different resistant mulberry genotypes treated with CK: Kangqing10, normal and control group (RN), Guisang12, normal and control group (SN); T1: Kangqing10-1 group (RS1), Guisang 12-1 group (SS1); T2: Kangqing 10-2 group (RS2), Guisang 12-2 group (SS2); T3: Kangqing 10-3 group (RS3), Guisang 12-3 group (SS3). Substrates: (A) amino acids, (B) amines, (C) polymers, (D) miscellaneous, (E) carboxylic acids, and (F) carbohydrates. The bars indicate the mean±SD (n = 3). Different letters indicate significant differences between treatments (P < 0.05).

Data concerning the Shannon evenness of microbial community structure are shown in Fig 8. The Shannon evenness of bacterial community structure was larger than that of the fungal community structure. The Shannon evenness of bacterial community structure in the resistant and susceptible genotypes was consistent with that of the fungal community structure in resistant and susceptible genotypes. The Shannon evenness values of the bacterial and fungal community structure of the RN rhizosphere genotypes were bothlower than those of the SN genotypes, but the Shannon evenness values of bacterial and fungal community structures of the RS1, RS2, and RS3 rhizosphere genotypes were larger than those of the SS1 and SS2 genotypes and significantly differed in SS1 (P < 0.05). The Shannon evenness values of the bacterial and fungal community structure of RN rhizosphere genotypes were both smaller than those of the RS1, RS2, and RS3 genotypes, but the Shannon evenness values of bacterial and fungal community structure of SN rhizosphere genotypes were both larger than those of the SS1, SS2, and SS3 genotypes, except for the SS3 fungal genotype. Additionally, the Shannon evenness of the bacterial community structure of the SN genotype was significantly different in SS1 and SS2, and the Shannon evenness values of the fungal community structure of the SN genotypes both displayed a significant difference in SS1 (P < 0.05). The Shannon evenness values of the bacterial and fungal community structure were both opposite to the Shannon index of soil microbial carbon source utilization, further indicating that the microbial community structure in susceptible genotypes is not as stable asthat in resistant genotypes which show some resistance to bacterial wilt.

**Fig 8.**
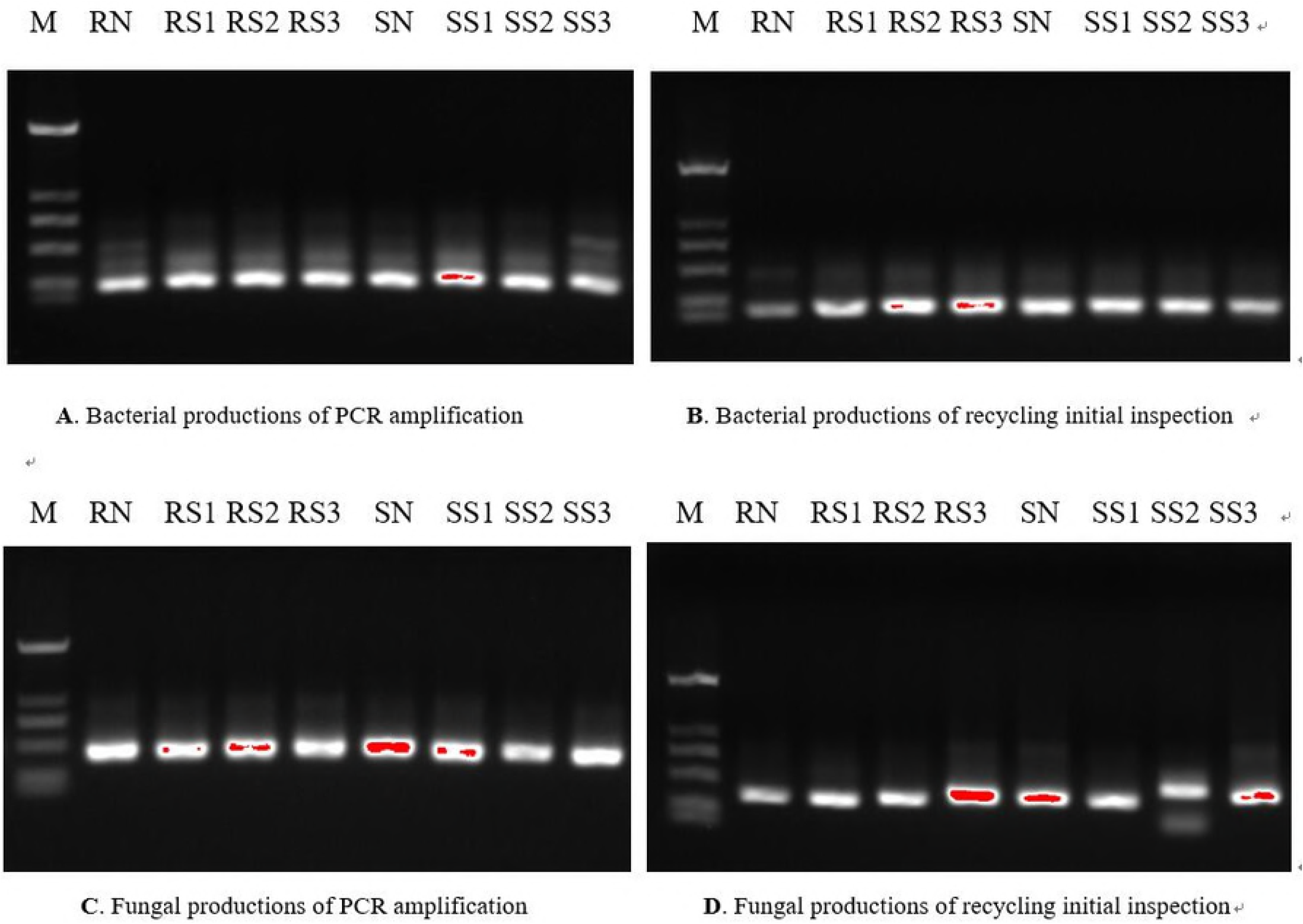
Comparison of the Shannon evenness (E) of bacterial and fungal community structure in soil rhizosphere with different resistant mulberry genotypes treated with CK Kangqing10, normal and control group (RN), Guisang12, normal and control group (SN); T1: Kangqing10-1 group (RS1), Guisang 12-1 group (SS1); T2: Kangqing 10-2 group (RS2), Guisang 12-2 group (SS2); T3: Kangqing 10-3 group (RS3), Guisang 12-3 group (SS3). Substrates: (A) amino acids, (B) amines, (C) polymers, (D) miscellaneous, (E) carboxylic acids, and (F) carbohydrates. The bars indicate the mean ± SD (n = 3). Different letters indicate significant differences between treatments (P < 0.05).

### Partial sequencing and NCBI alignment

NCBI alignments of bacteria and fungi are shown in Table 3 and 4, respectively. There were more bacterial species than fungal species in the soil rhizosphere with different resistant mulberry genotypes. Ralstonia solanacearum was found in the rhizosphere of resistant and susceptible genotypes, indicating that bacterial wilt likely occurred because of the destruction of the bacterial community structure in mulberry rhizosphere soil by R. solanacearum.

**Table 3.**
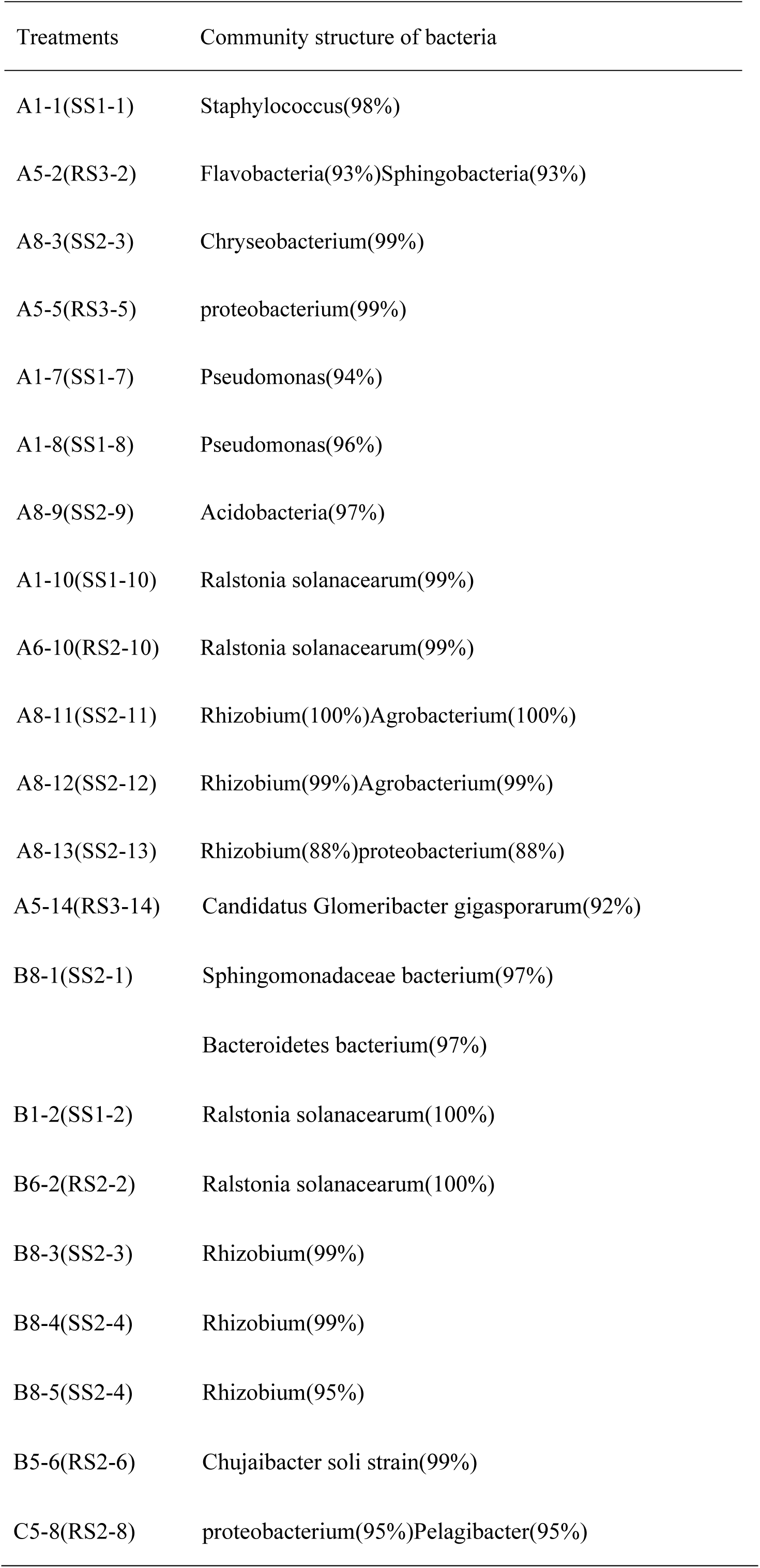
NCBI alignment of bacteriarandomly recovered from the bacterial DGGE profiles

**Table 4.**
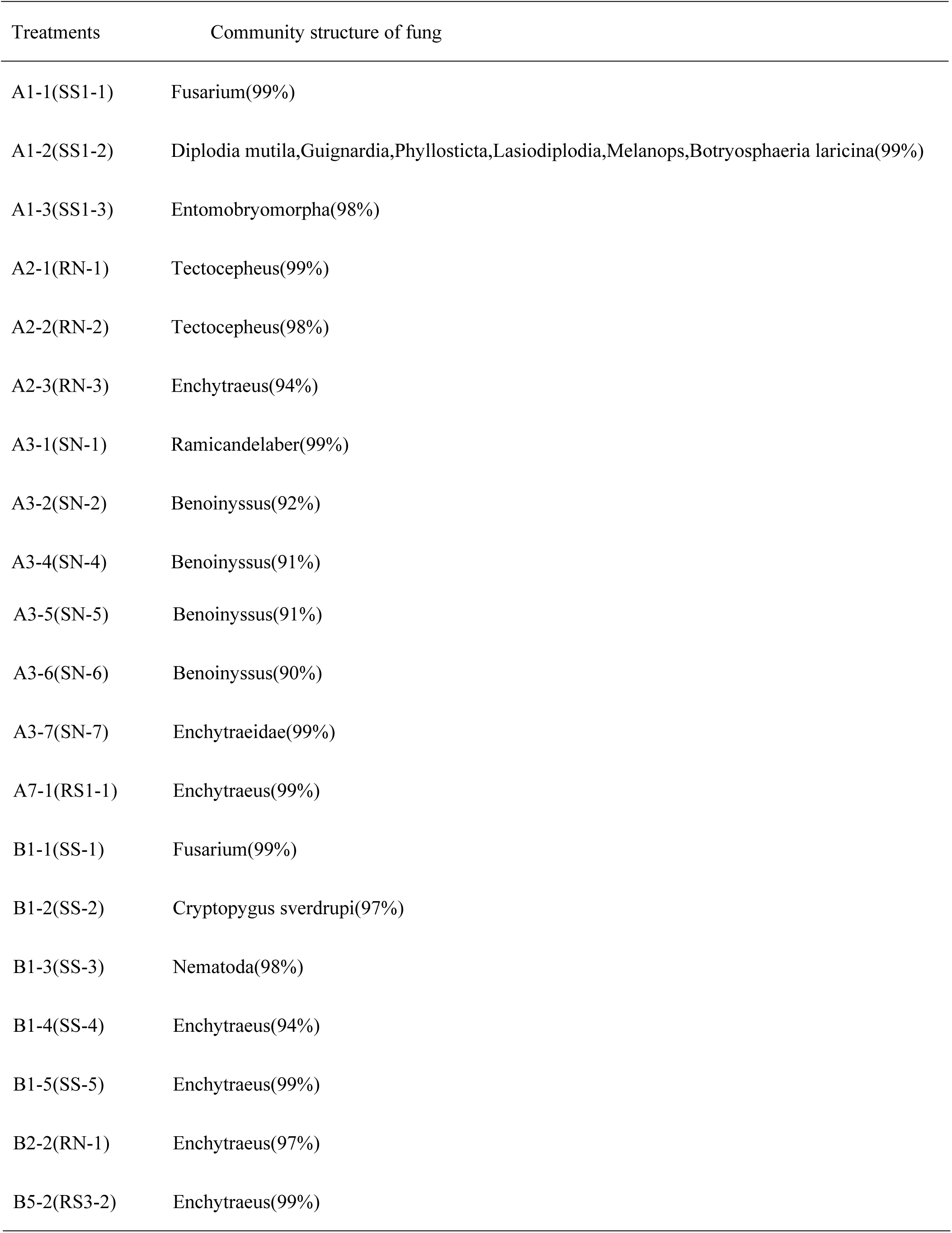
NCBI alignment of fungal randomly recovered from the fungal DGGE profiles

| Treatments | Community structure of fung |
| --- | --- |
| A1-1(SS1-1) | Fusarium(99%) |
| A1-2(SS1-2) | Diplodia mutila,Guignardia,Phyllosticta,Lasiodiplodia,Melanops,Botryosphaeria laricina(99%) |
| A1-3(SS1-3) | Entomobryomorpha(98%) |
| A2-1(RN-1) | Tectocephus(99%) |
| A2-2(RN-2) | Tectocephus(98%) |
| A2-3(RN-3) | Enchytraeus(94%) |
| A3-1(SN-1) | Ramicandelaber(99%) |
| A3-2(SN-2) | Benoinyssus(92%) |
| A3-4(SN-4) | Benoinyssus(91%) |
| A3-5(SN-5) | Benoinyssus(91%) |
| A3-6(SN-6) | Benoinyssus(90%) |
| A3-7(SN-7) | Enchytraeidae(99%) |
| A7-1(RS1-1) | Enchytraeus(99%) |
| B1-1(SS-1) | Fusarium(99%) |
| B1-2(SS-2) | Cryptopygus sverdrupi(97%) |
| B1-3(SS-3) | Nematoda(98%) |
| B1-4(SS-4) | Enchytraeus(94%) |
| B1-5(SS-5) | Enchytraeus(99%) |
| B2-2(RN-1) | Enchytraeus(97%) |
| B5-2(RS3-2) | Enchytraeus(99%) |

### Correlation regression analysis between soil microbial community structure and soil nutrients

Correlation regression analysis data between bacterial and fungal community structure and soil nutrientsare presented in Table 5 and 6, respectively. The bacterial and fungal community structures of mulberry resistant rhizosphere genotypes were significantly correlated with soil organic matter (R=0.9697 and 0.9666, respectively) and alkaline nitrogen (R=0.9519 and 0.9759, respectively), but displayed a weak correlation with available soil phosphorus and potassium. The bacterial and fungal community structures of the mulberry susceptible genotypes in rhizosphere were also significantly correlated with soil organic matter and alkaline nitrogen, except forbacterial alkaline nitrogen. This may be because of thehigh humidity, temperature, and rainfall in Southern China. More frequent precipitation and higher humidity accelerate soil decomposition and mineralization. As a result, the occurrence of mulberry bacterial wilt may be related to the loss of soil nutrients, particularly soil organic matter and nitrogen. These losses may cause an imbalance in the soil microbial community structure. In contrast, the correlations between the bacterial and fungal community structures of mulberry resistant genotypes in the rhizosphere and soil organic matter (R=0.9697 and 0.9666, respectively) and alkaline nitrogen (R=0.9519 and0.9759, respectively)were greater than those between the bacterial and fungal community structures of mulberry susceptible genotypes in the rhizosphere and soil organic matter (R= 0.8808 and 0.8993, respectively) and alkaline nitrogen (R=0.0745 and 0.6899, respectively). These results indicate that the loss of soil organic matter and nitrogen has a lower impact on resistant genotypes than on susceptible genotypes. Thus, the bacterial and fungal community structures in resistant genotypes aremore stable than in susceptible genotypes.

**Table 5.**
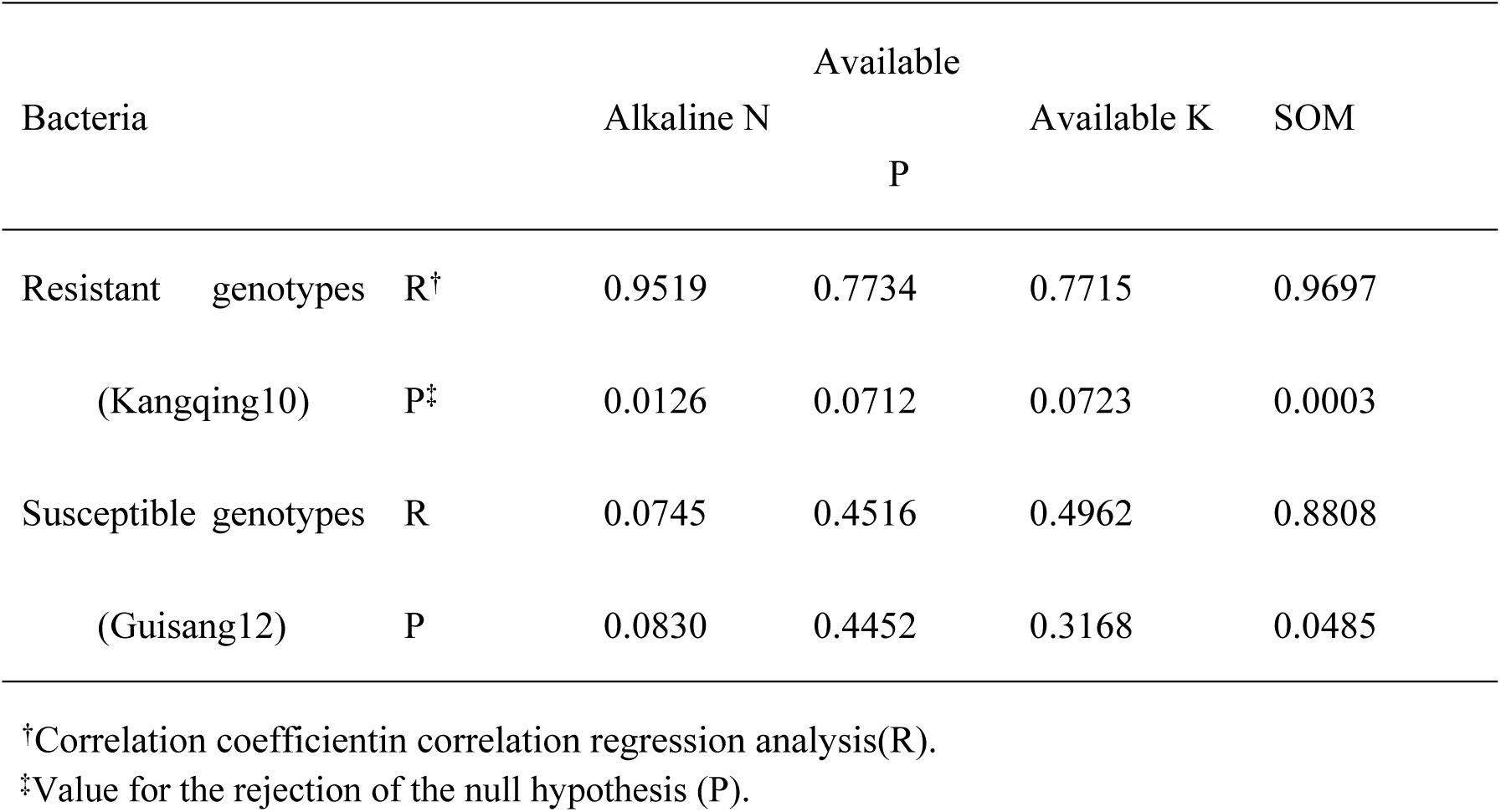
Correlation regression analysis of that between soil nutrients and soil bacterial community structure in rhizosphere treated with resistant genotypes (Kangqing10) and susceptible genotypes (Guisang12), including alkaline N,alkaline P, alkaline K, and soil organic matter (SOM).

**Table 6.**
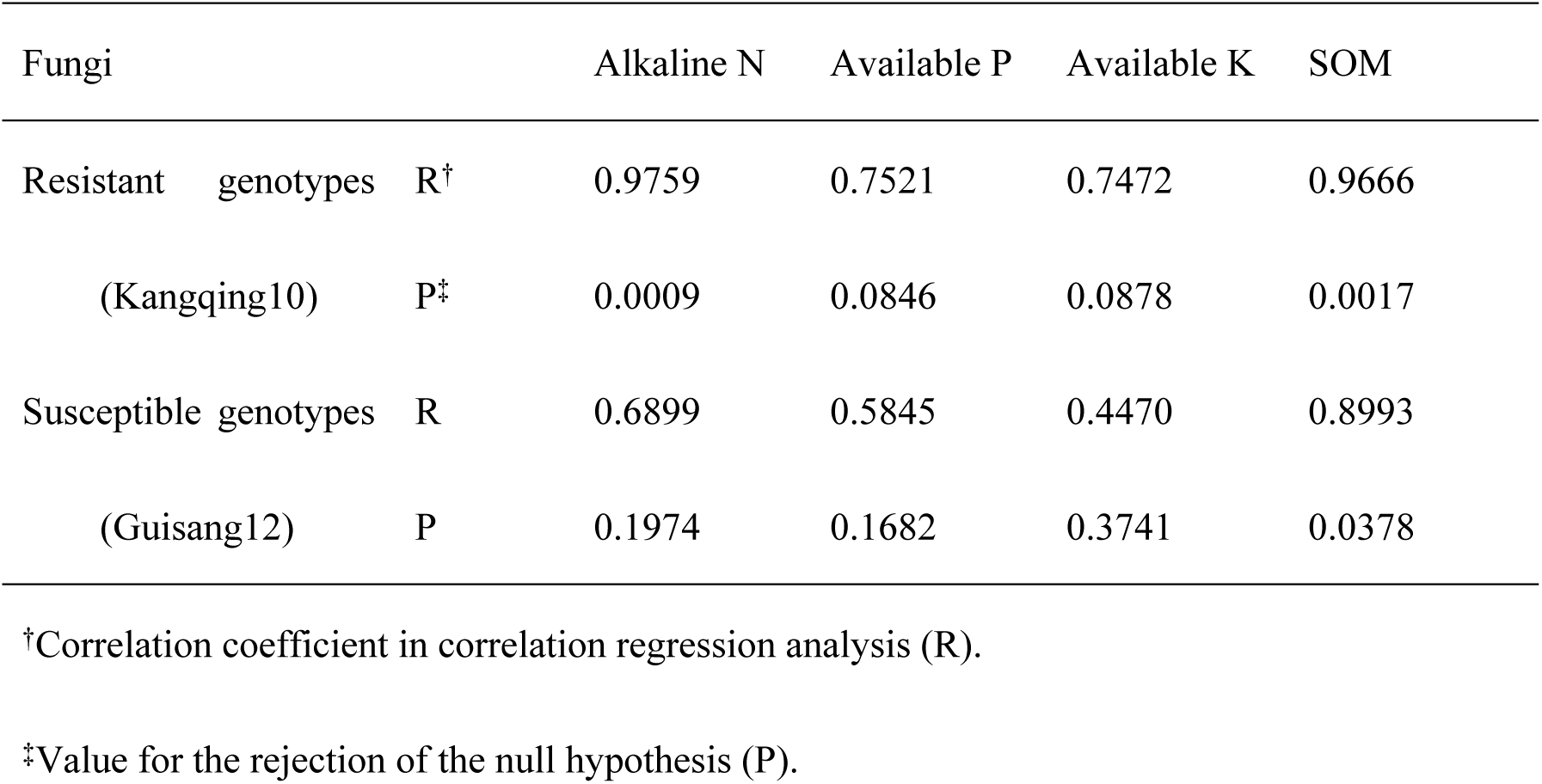
Correlation regression analysis of that between soil nutrients and soil fungal community structure in rhizosphere treated with resistant genotypes (Kangqing10) and susceptible genotypes (Guisang12), both including alkaline N,alkaline P, alkaline K and soil organic matter (SOM)

| Fungi |  |  | Alkaline N | Available P | Available K | SOM |
| --- | --- | --- | --- | --- | --- | --- |
| Resistant | genotypes | R<sup>†</sup> | 0.9759 | 0.7521 | 0.7472 | 0.9666 |
|  | (Kangqing10) | P<sup>‡</sup> | 0.0009 | 0.0846 | 0.0878 | 0.0017 |
| Susceptible | genotypes | R | 0.6899 | 0.5845 | 0.4470 | 0.8993 |
|  | (Guisang12) | P | 0.1974 | 0.1682 | 0.3741 | 0.0378 |

†Correlation coefficient in correlation regression analysis (R).
‡Value for the rejection of the null hypothesis (P).

## Discussion

The health of plants is affected by interactions among plants, the soil environment, and soil microorganisms. Changes in one or more oftheelements can lead to plant diseases. As a dynamic part in the plant-soil system,the homeostasis of soil microorganisms is extremely important for controlling bacterial wilt and other soil-borne diseases [37-42]. Resistant mulberry genotypes are more stable than susceptible mulberry genotypes and exhibit better protection against bacterial wilt. Thus, the incidence of bacteria wilt is relatively low. Recent studies have reported that eggplant, tobacco, and pepper plants with a resistant genotypehave a significantly reduced incidence of bacterial wilt [43-47]. The present study was conducted to explorethe relationship between the occurrence of bacterial wilt and abundance and community structure diversity of soi microorganisms with plants of different genotypes. The genotypes included resistant normal mulberry genotype plants(RN), susceptible normal mulberry genotype plants (SN), resistant sickly mulberry genotype plants(RS), and susceptible sickly mulberry genotype plants (SS). This is the first study to show similar changes in the abundance and carbon source utilization of soil microorganisms in mulberry genotypes resistant or susceptible to bacterial wilt. Additionally, changes in microbial community structures were significantly correlated with soil organic matterand alkaline nitrogen. For normal plants, the abundance of bacteria in resistant mulberry genotypes was higher than that in susceptible mulberry genotypes. For sickly plants, the abundance of bacteria in resistant mulberry genotypes was lower than that in susceptible mulberry genotypes. Average well color development values and the Shannon index of carbon source utilization revealed significantly better utilization of carbon sources by resistant normal mulberry rhizosphere genotypes than by susceptible normal mulberry genotypes. In contrast, carbon sources utilization in the RS rhizosphere genotypeswas poorer than that in the SS genotypes. These observations are related to the abundance of soil microorganisms. Irikiin Y [48] has suggested that the occurrence of tomato bacterial wilt is related to the pattern of carbon source utilization by rhizosphere soil bacteria. The present PCR-DGGE examination revealed that the microbial community structures of RN rhizosphere genotypes were not as rich as those of SN genotypes. However, the microbial community structure of the RS rhizosphere genotypes was richer than that ofthe SS genotypes. Finally, regression correlation analysis of the soil microbial community structure and soil nutrients revealed that loss of soil organic matter and nitrogen may have a greater negative impact on susceptible mulberry genotype plants than on resistant mulberry genotype plants, and the microbial community structures in resistant mulberry genotype plants are more stable than those in susceptible mulberry genotype plants.

The collective results of previous [49, 50] and the present study suggest that the occurrence of mulberry bacterial wilt is related to the loss of soil nutrients, particularly organic matter and nitrogen, which can disrupt the soil microbial community structure. However, these losses have a lower impact on resistant genotype plants than on susceptible genotype plants. Therefore, resistant genotype plants have some resistance to bacterial wilt. A previous study [51] indicated that the rate of utilization of soil nutrientsis greater in resistant genotype plants than in susceptible genotype plants. In the current study, carbon utilization in resistant genotype plants was better than that insusceptible genotype plants. Our empirical data indicate that susceptible genotype plants are more dependent on soil nutrients than are resistant genotype plants.

Our results will be useful for developing methods to prevent and control bacteria wilt involving the application oforganic fertilizer and nitrogen-containing fertilizer. These fertilizerscan revitalize nutritionally depleted soil. Mulberry trees display growth vitality that can naturally prevent and control bacteria wilt naturally. Further studies in our lab will focus on the micro-ecological mechanisms of mulberry bacterial wilt, emphasizing the microbes in the surrounding rhizosphere by choosing resistant genotype varieties and appropriately applying organic fertilizer or nitrogen fertilizer.

## Acknowledgements

The project was financially supported by the National Natural Science Foundation of China (No. 31770660 and 31300518) and the Earmarked Fund for Modern Agro-industry Technology Research System (CARS-22).

---

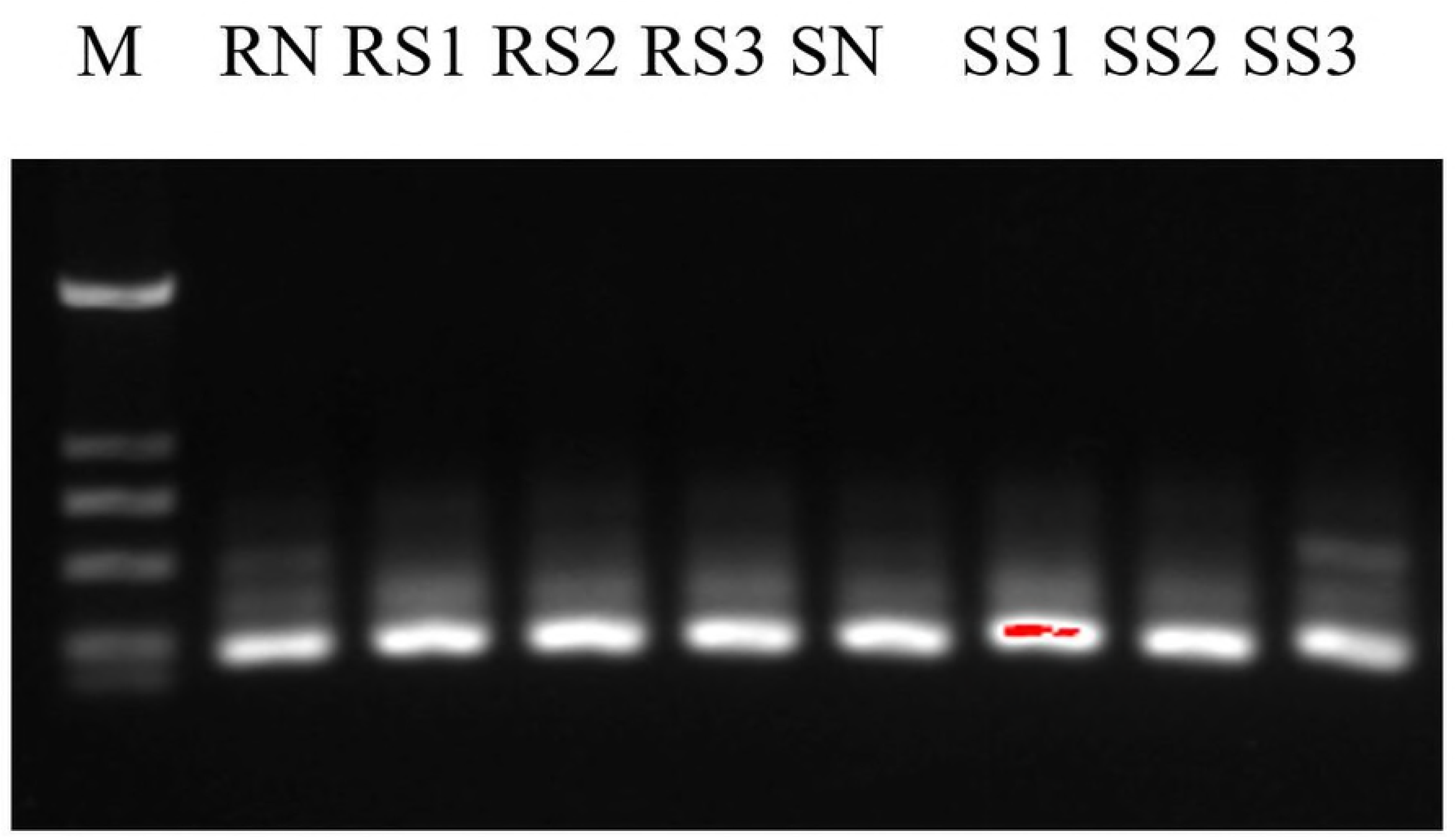

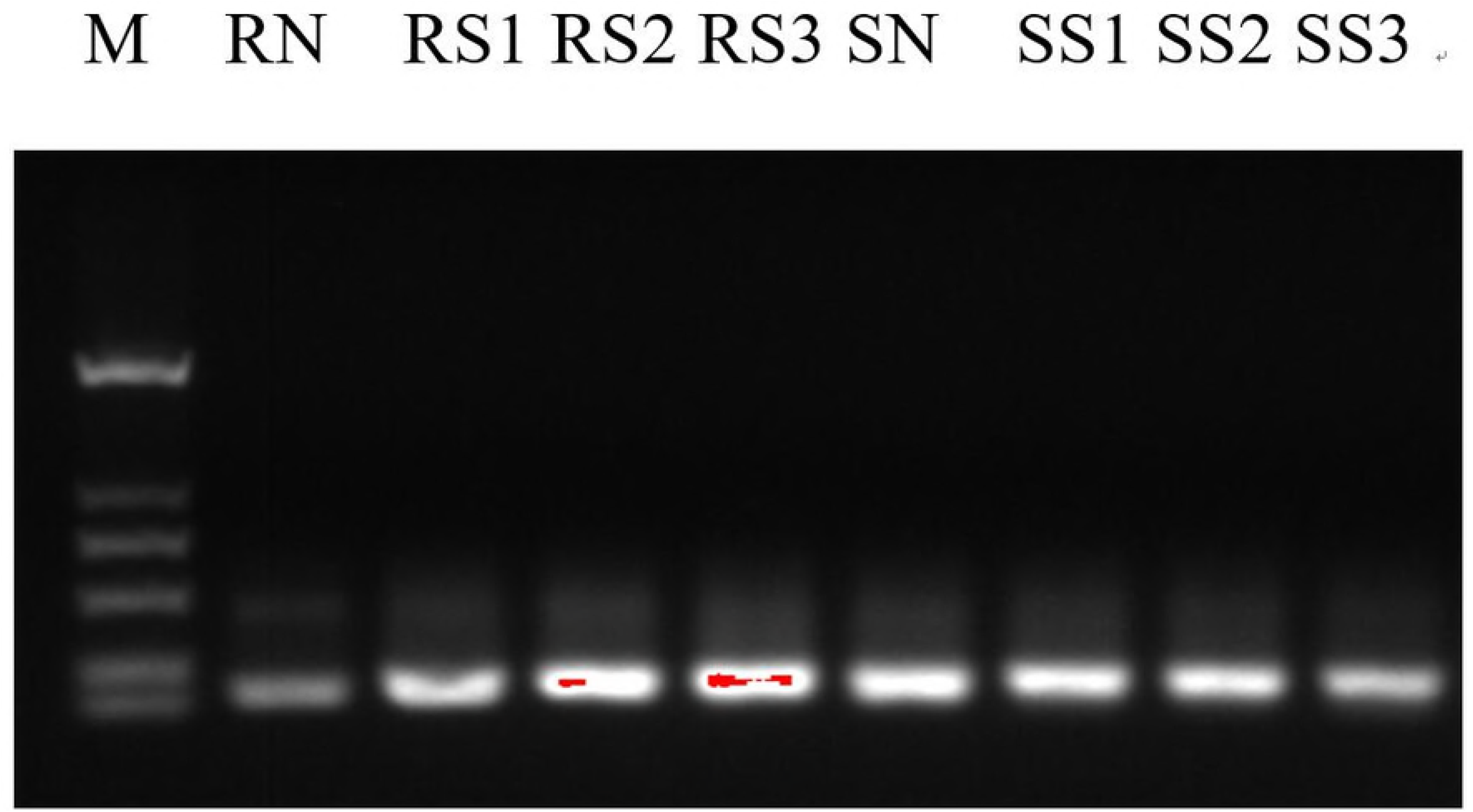

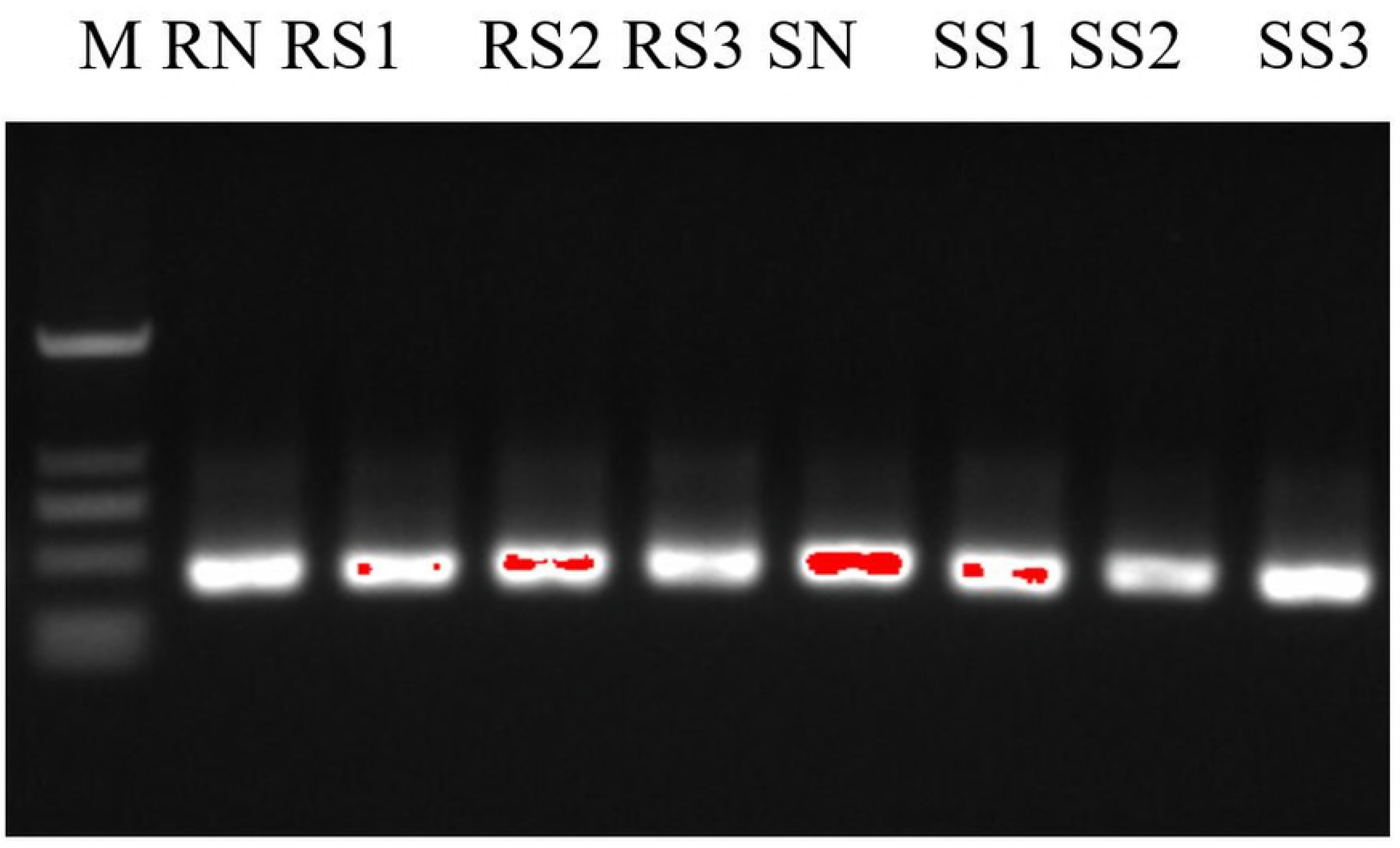

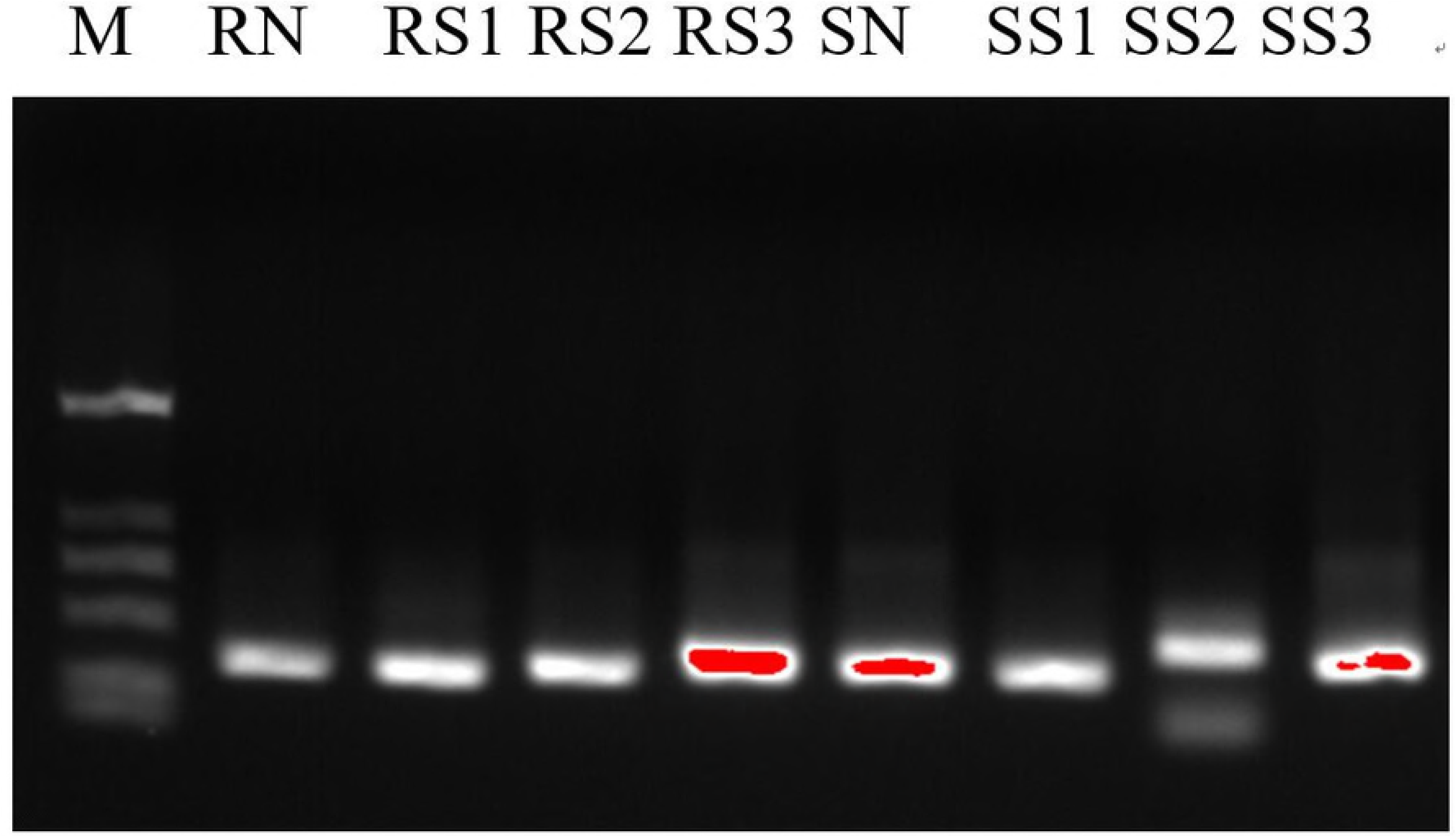

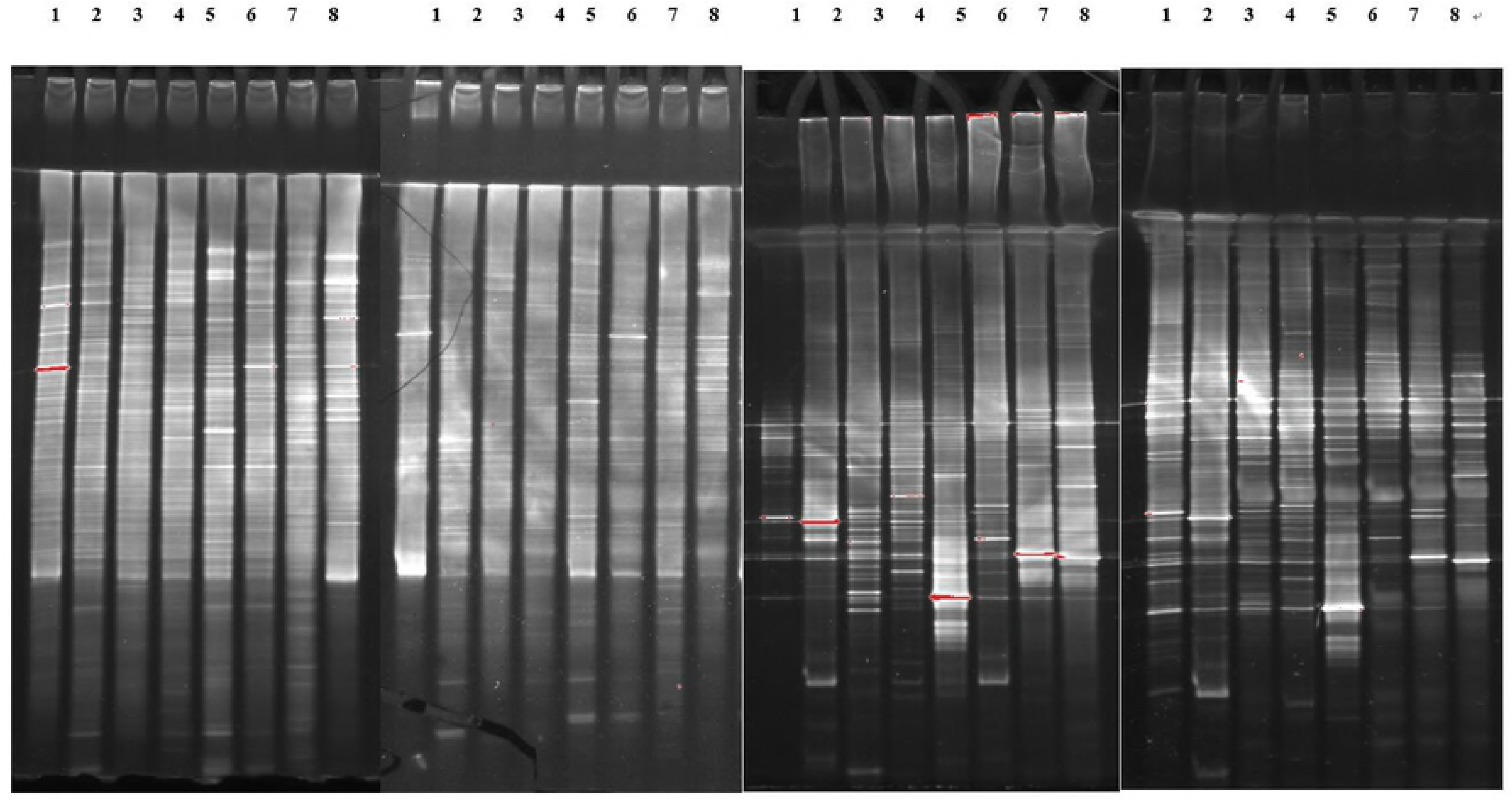

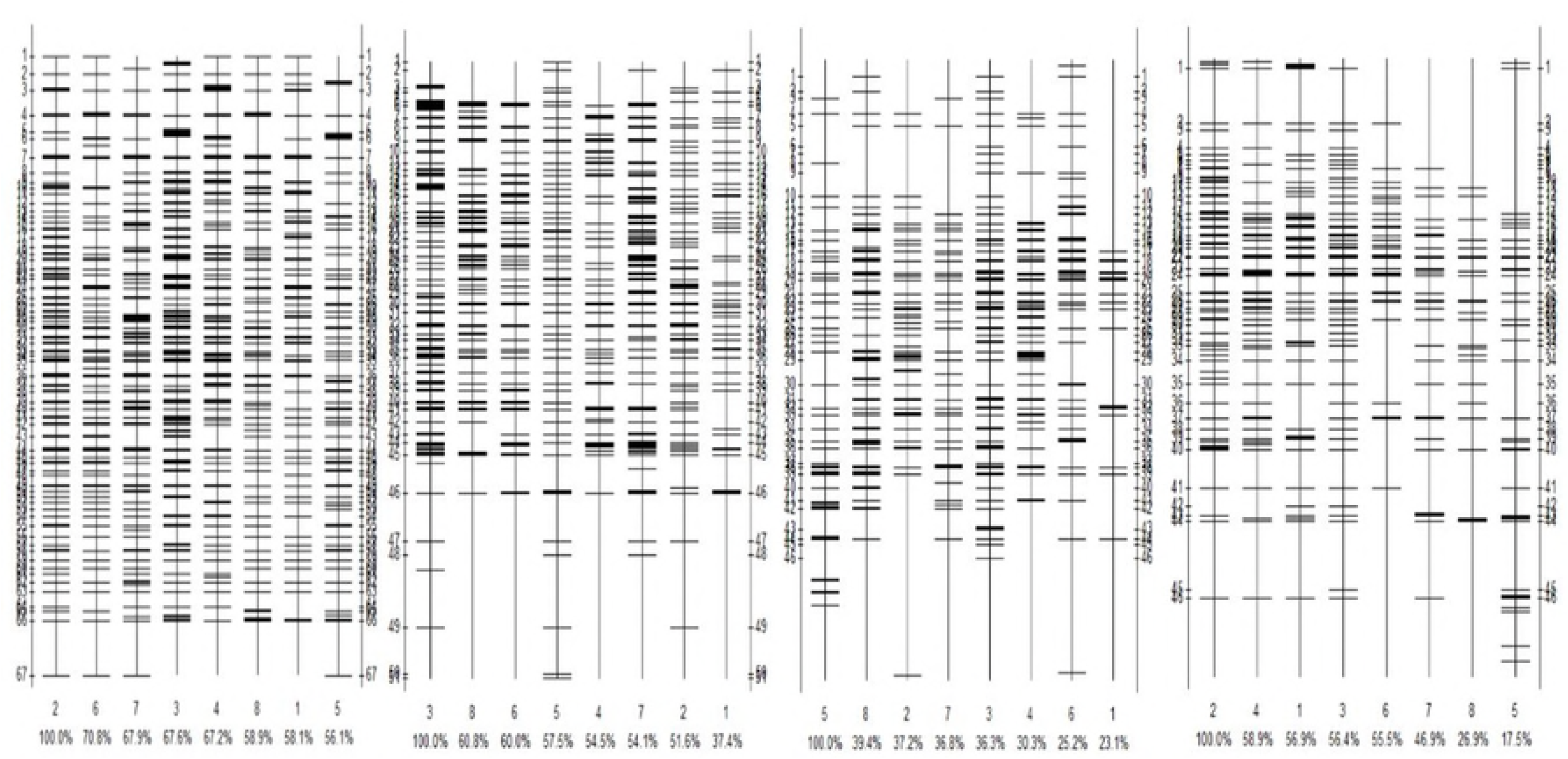

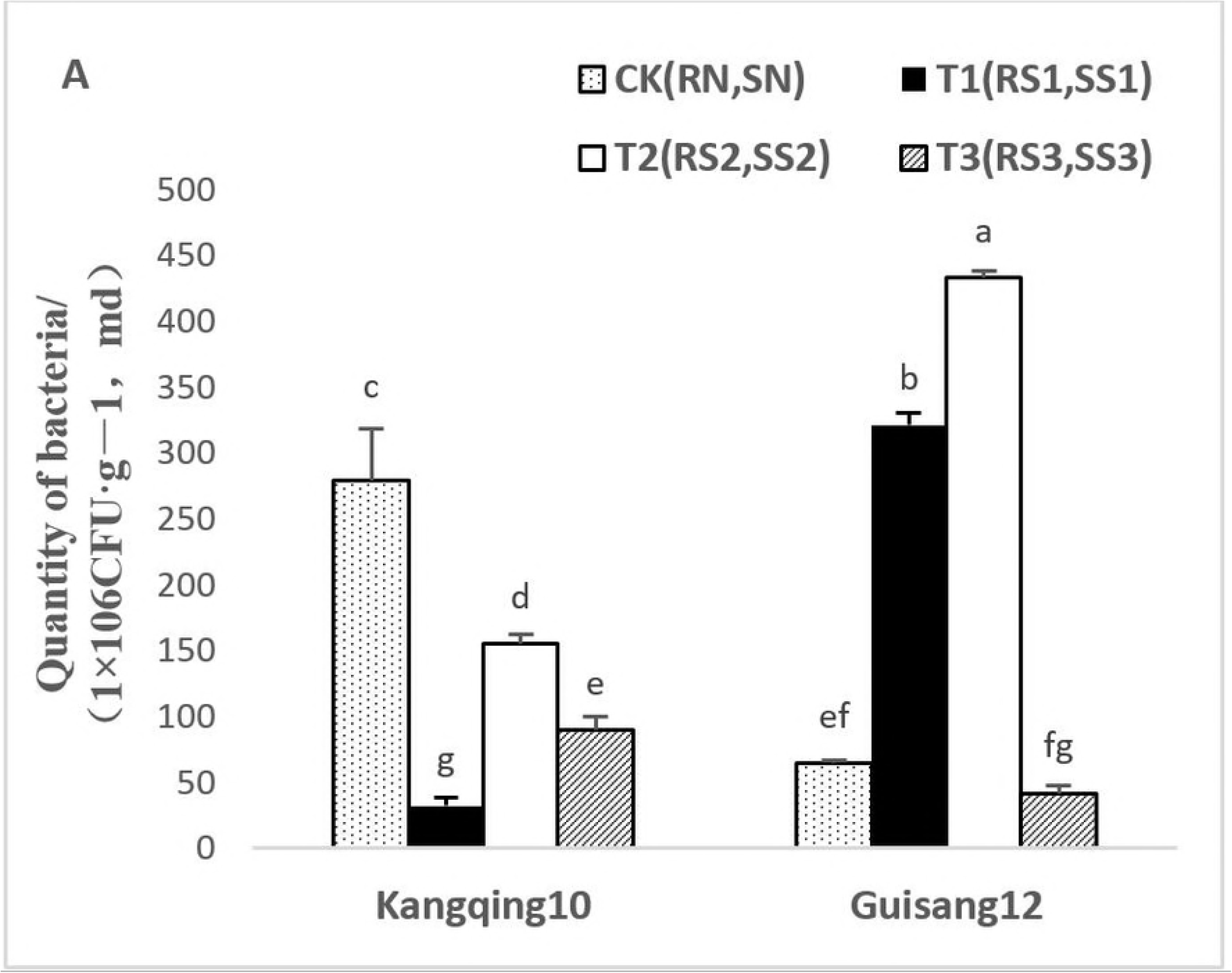

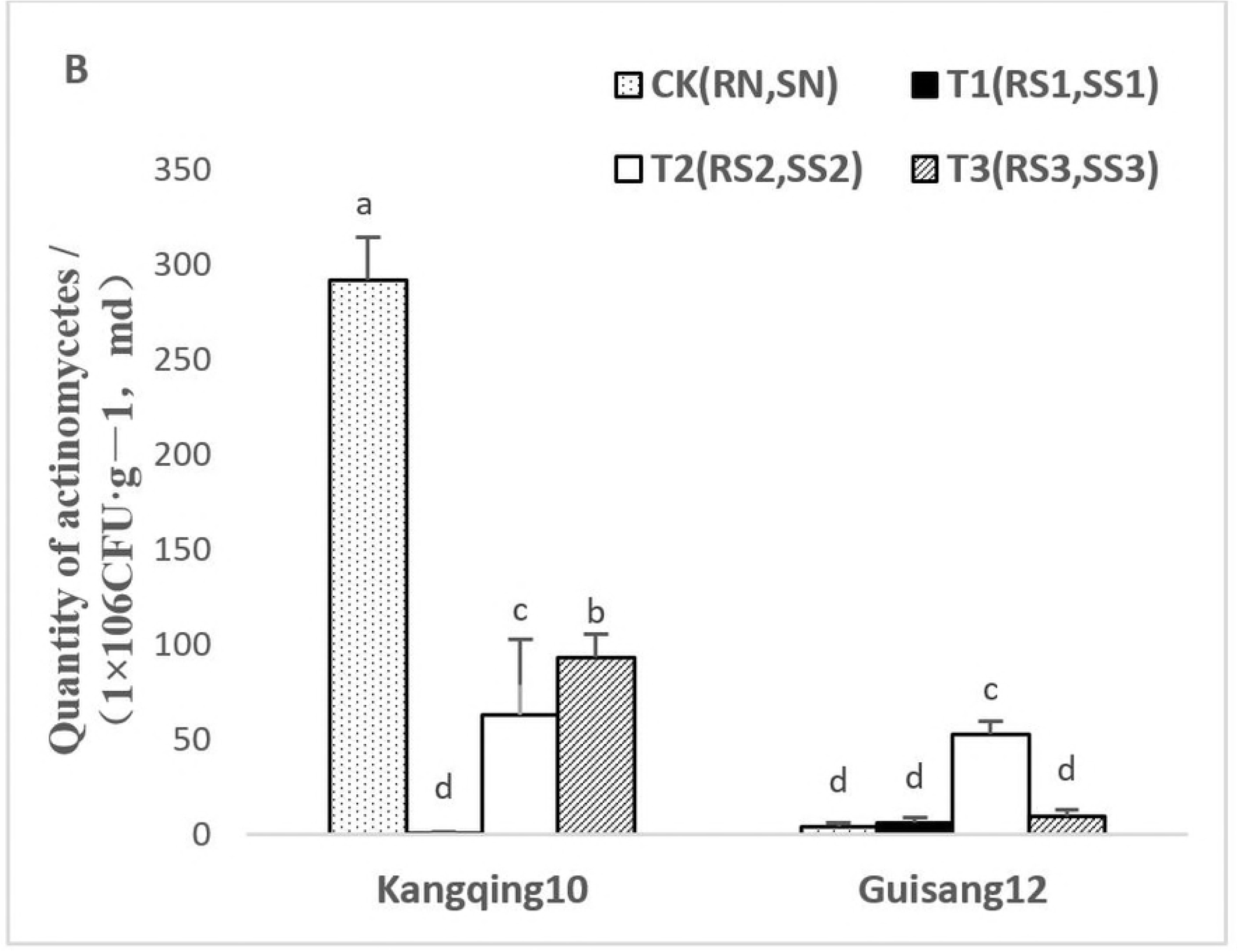

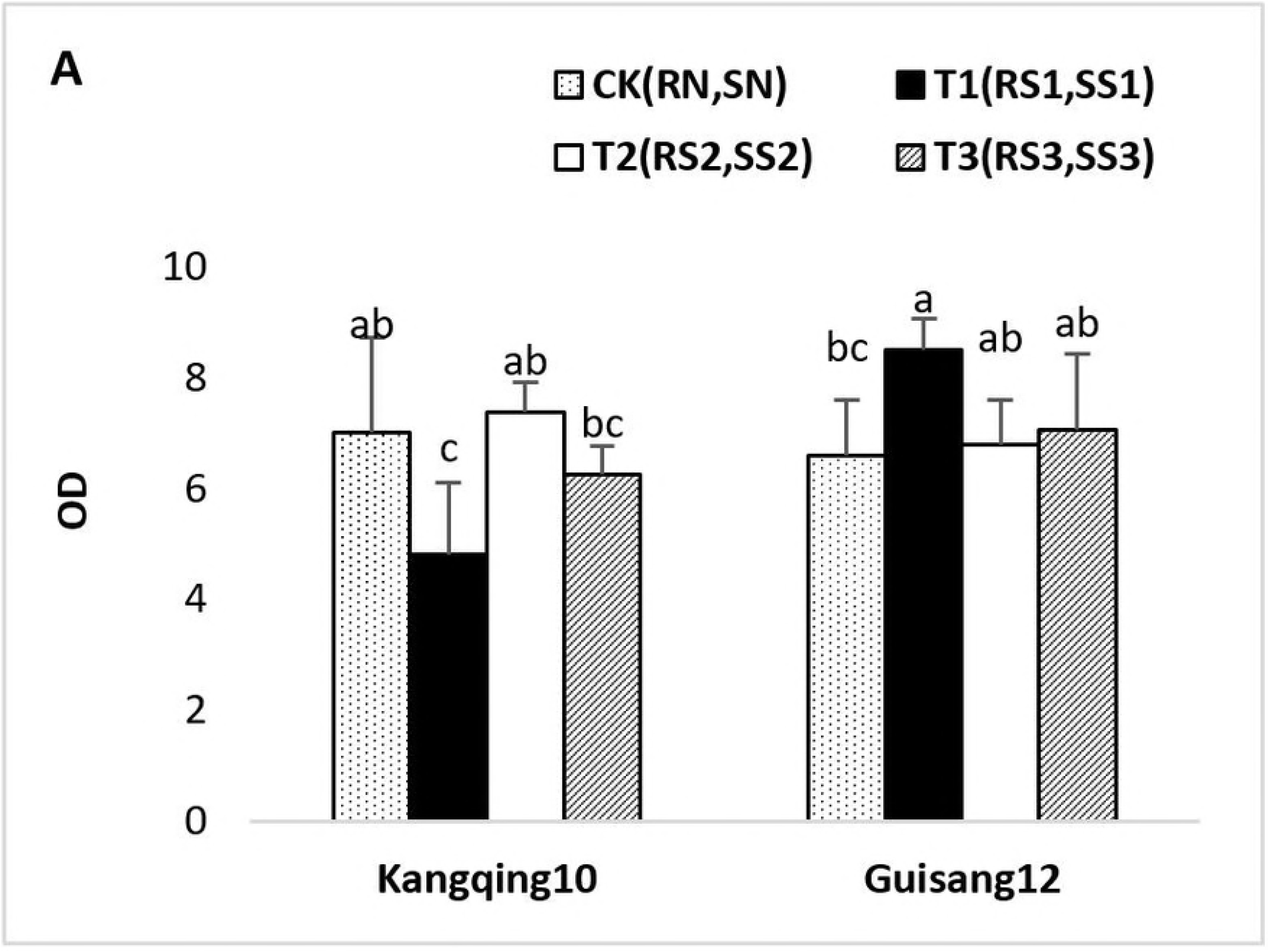

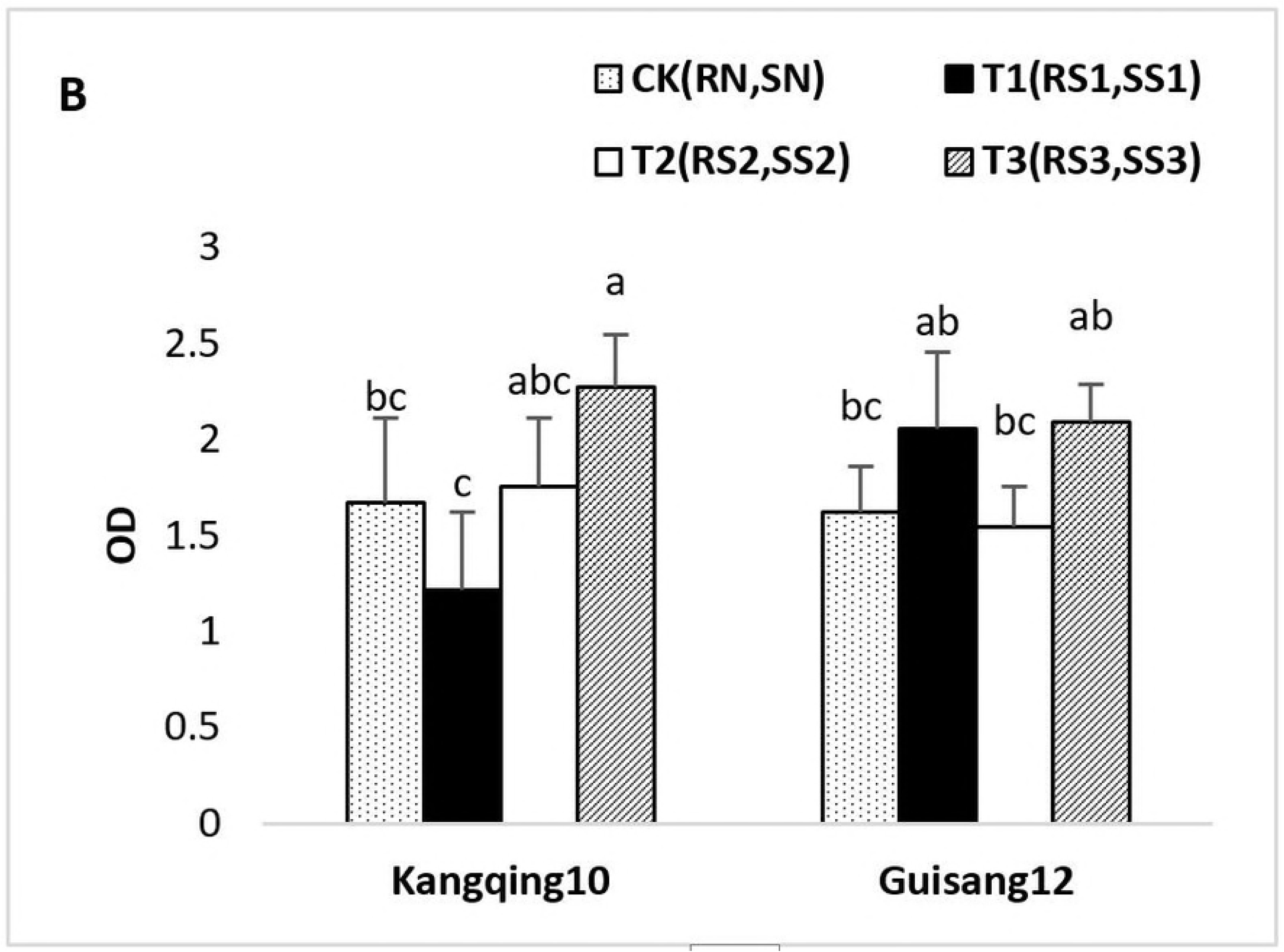

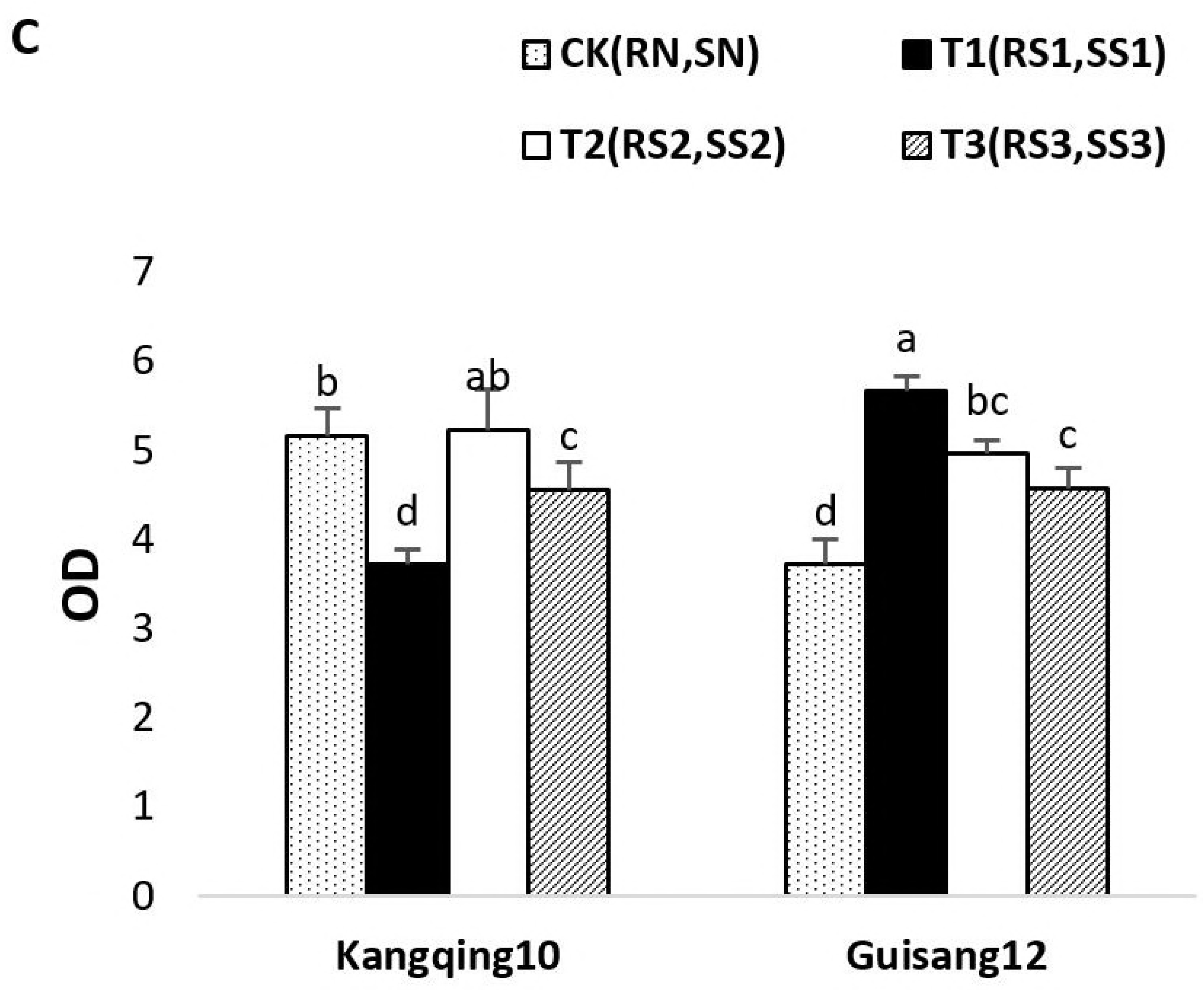

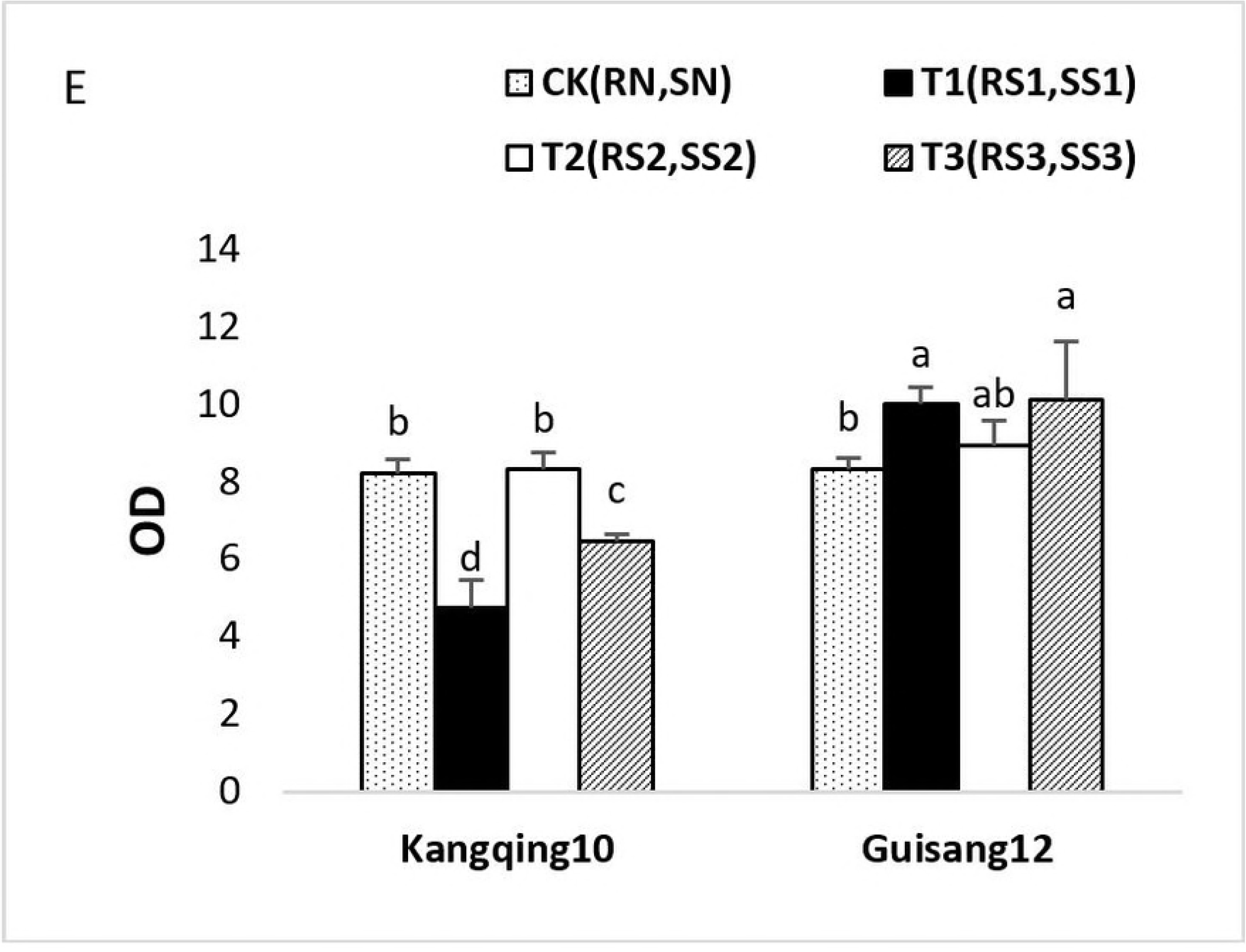

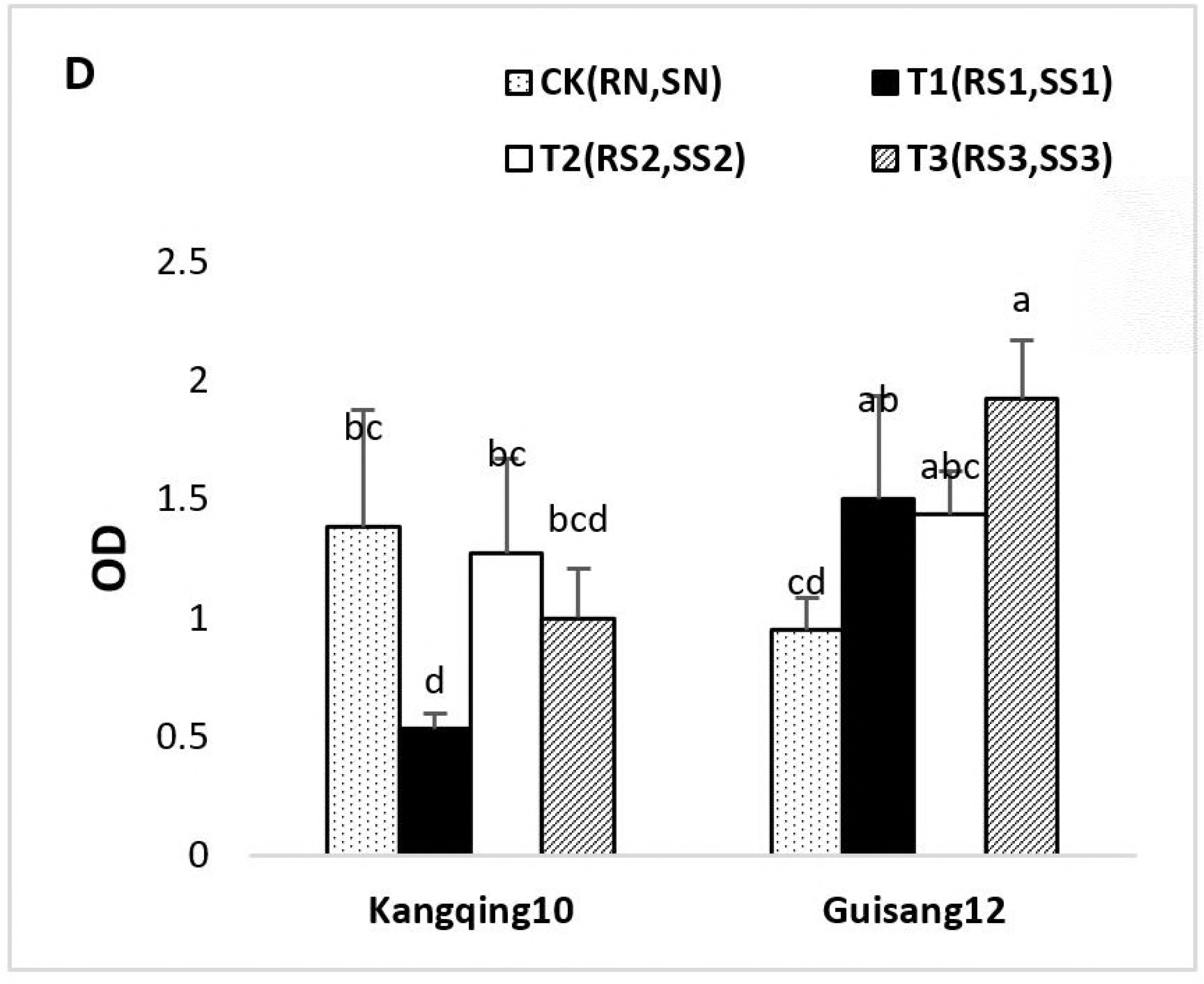

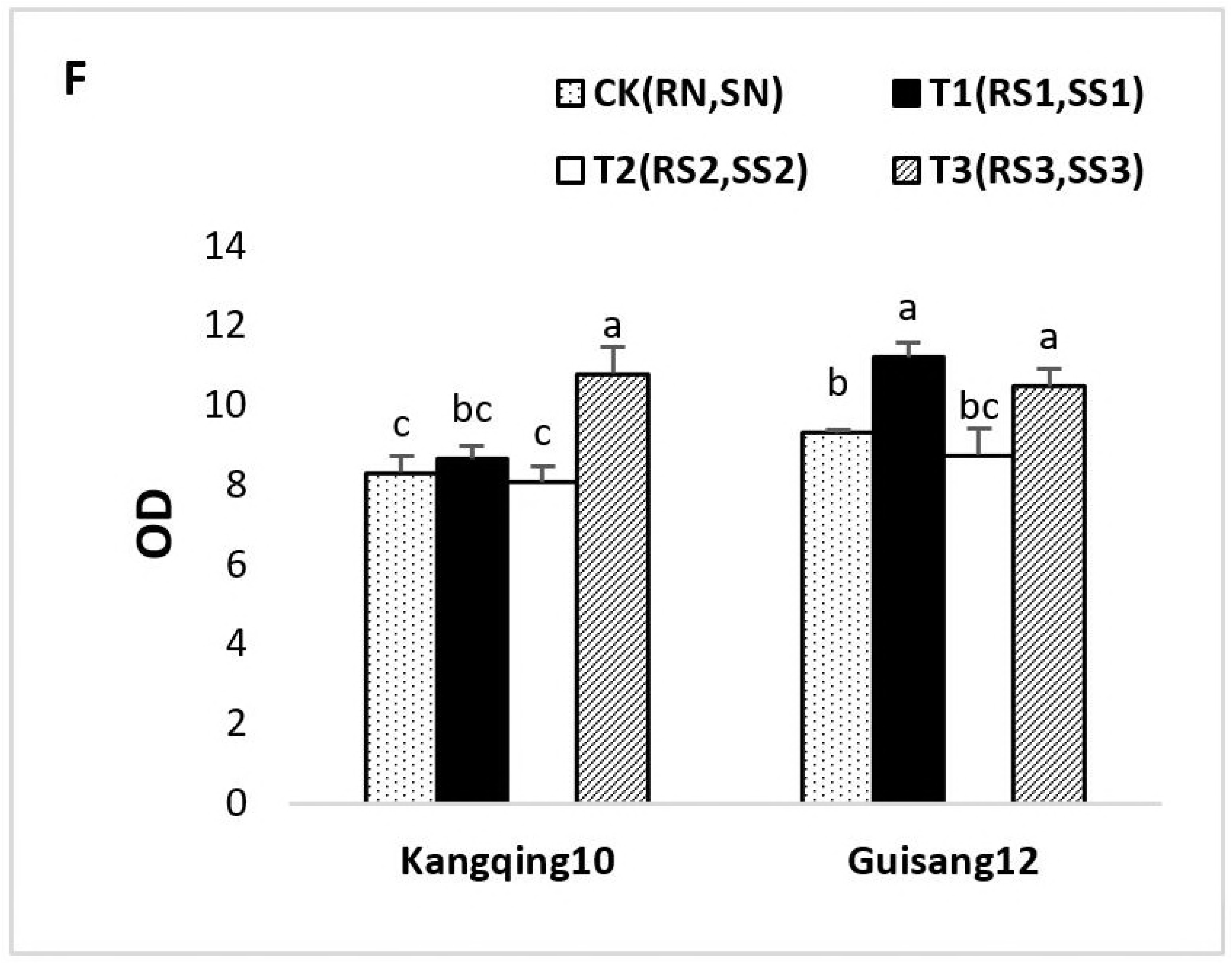

## References

1. Hayward AC. Biology and Epidemiology of Bacterial Wilt Caused by Pseudomonas Solanacearum. Annurevphytopathol. 2003;29(1):65–87.

2. Kariko EK, Murimi ZK, Owuor PO, Maingi JM. Bacterial wilt, a challenge in Solanaceous crops production at Kenyan highlands and lowlands. World Journal of Research and Review. 2016;3: 2455–3956‥

3. Couëdel A, Alletto L, Kirkegaard J, Justes É. Crucifer glucosinolate production in legume-crucifer cover crop mixtures. European Journal of Agronomy. 2018;96:22-33.

4. Xiao Y, Liu X, Meng D, Tao J, Gu Y, Yin H, et al. The role of soil bacterial community during winter fallow period in the incidence of tobacco bacterial wilt disease. Applied Microbiology & Biotechnology. 2018;102(5):2399–412.

5. Rand F V. Dissemination of bacterial wilt of cucurbits. Journal of Agronomic Research. 1915;6(4):417–434.

6. Zhao Q, Wu J, Zhang L, Xu L, Yan C, Gong Z. Identification and characterization of Cucurbita gummy stem blight fungi in Northeast China. Journal of Phytopathology. 2018;166(6).

7. Wang GF, Xie GL, Zhu B, Huang JS, Liu B, Kawicha P, et al. Identification and characterization of the Enterobacter complex causing mulberry (Morus alba) wilt disease in China. European Journal of Plant Pathology. 2010;126(4):465–78.

8. Mahadevakumar S, Chandana C, Deepika YS, Sumashri KS, Yadav V, Janardhana GR. Pathological studies on the southern blight of China aster (Callistephus chinensis) caused by Sclerotium rolfsii. European Journal of Plant Pathology. 2018(3):1–7.

9. Qiu J, Xu L, Qian Y, Zeng Y, Hou Y, Zhang N, et al. An Investigation on Rhizospheric Bacterial Community and Soil Physical and Chemical Properties of Different Mulberry Varieties. Science of Sericulture. 2017;88(8).

10. Ashaduzzaman SM, Israt ZM, Li CF, Dai CC. Endophytic fungusPhomopsis liquidambariand different doses of N-fertilizer alter microbial community structure and function in rhizosphere of rice. Scientific Reports. 2016;6:32270.

11. Ghose L, Neela FA, Chakravorty TC, Ali MR, Alam MS. Incidence of leaf blight disease of mulberry plant and assessment of changes in amino acids and photosynthetic pigments of infected leaf. Plant Pathology Journal. 2010;9(3).

12. Cen ZL, Ouyang QF, Xie L, Chen BS, Zhu FR, Lin Q, et al. Isolation and Identification of Pathogenic Bacteria from Mulberry Bacterial Wilt in Guangxi. Southwest China Journal of Agricultural Sciences. 2013;26:1054–1057.

13. Zhang JJ, Liu H, Xia LI. Effects of combined application of different organic fertilizers on the quality and the incidence of the soil-borne diseases of greenhouse strawberry. Journal of Anhui Agricultural University. 2013;1301–1305.

14. Wang F, Yuan T, Shoukuan GU, Wang Z. Effects of organic and inorganic slow-release compound fertilizers on microbial biomass carbon and nitrogen,and microbial community structure in soil. Acta Ecologica Sinica. 2016;36(7).

15. Illera-Vives M, Labandeira SS, Loureiro LI, López-Mosquera ME. Agronomic assessment of a compost consisting of seaweed and fish waste as an organic fertilizer for organic potato crops. Journal of Applied Phycology. 2017;29(3):1663–71.

16. Guo J, Liu H, Li S. Biocontrol agent against soil-borne diseases. EP2255660 A1; 2012.

17. Ayana G, Fininsa C, Ahmed S, Wydra K. Effects of Soil Amendment on Bacterial Wilt Caused by Ralstonia Solanacerum and Tomato Yields in Ethiopia. Journal of Plant Protection Research. 2011;51(1):72–6.

18. Ayana G, Fininsa C, Ahmed S, Wydra K. Effects of Soil Amendment on Bacterial Wilt Caused by Ralstonia Solanacerum and Tomato Yields in Ethiopia. Journal of Plant Protection Research. 2011;51(1).

19. Niu J, Chao J, Xiao Y, Chen W, Zhang C, Liu X, et al. Insight into the effects of different cropping systems on soil bacterial community and tobacco bacterial wilt rate. J Basic Microbiol. 2017;57(1):3–11.

20. Ji X. Biological control against bacterialwilt and colonizationof mulberrybyan endophytic Bacillussubtilis strain. FEMS Microbiology Ecology. 2008;565–573.

21. Kazuo M. Soil sampling device and method for collecting an undisturbed soil sample. DE3125239. 1982.

22. Riley D, Barber SA. Bicarbonate Accumulation and pH Changes at the Soybean (Glycine max (L.) Merr.) Root-Soil Interface 1. Soil Science Society of America Journal. 1969;33(6):905–8.

23. Bao SD. Soil and agricultural chemistry analysis. China Agriculture Press. 2000;30–34.

24. Blakemore LC. Methods for chemical analysis of soils. New Zealand Soil Bureau Scientific Report. 1987;80:19–22.

25. Pinta M. Agricultural Applications of Flame Photometry: Macmillan Education UK; 1970.

26. Nelson DW, Sommers LE. Total carbon, organic carbon and organic matter, in: Methods of Soil Analysis Part 2. Chemical and Microbial Properties. 1982; 539–579.

27. Hirte WF. [The use of dilution plate method for the determination of soil microflora. 2. The qualitative demonstration of bacteria and actinomycetes]. Zentralblatt für Bakteriologie, Parasitenkunde, Infektionskrankheiten und Hygiene Zweite naturwissenschaftliche Abt: Allgemeine, landwirtschaftliche und technische Mikrobiologie. 1969;123(2):167.

28. You J, Chen MB, Fang SL, Liu QS. Selection of Isolation Media of Microbes in the Fermentation Pit of Chinese Strong Aromatic Spirits. Liquor Making. 2009; 262(3).

29. Garland JL, Mills AL. Classification and characterization of heterotrophic microbial communities on the basis of patterns of community-level sole-carbon-source utilization. Appl Environ Microbiol. 1991;57(8):2351–9.

30. Garland JL. Analytical approaches to the characterization of samples of microbial communities using patterns of potential C source utilization. Soil Biology & Biochemistry. 1996;28(2):213–21.

31. Zak JC, Willig MR, Moorhead DL, Wildman HG. Functional diversity of microbial communities: A quantitative approach. Soil Biology & Biochemistry. 1994;26(9):1101–8.

32. Staddon WJ, Duchesne LC, Trevors JT. Microbial Diversity and Community Structure of Postdisturbance Forest Soils as Determined by Sole-Carbon-Source Utilization Patterns. Microbial Ecology. 1997;34(2):125–30.

33. Tresse O, Mounien F, Lévesque MJ, Guiot S. Comparison of the microbial population dynamics and phylogenetic characterization of a CANOXIS reactor and a UASB reactor degrading trichloroethene. Journal of Applied Microbiology. 2005;98(2):440–9.

34. Bottos EM, Vincent WF, Greer CW, Whyte LG. Prokaryotic diversity of arctic ice shelf microbial mats. Environmental Microbiology. 2008;10(4):950–66.

35. Wang HY, Gao YB, Fan QW, Xu Y. Characterization and comparison of microbial community of different typical Chinese liquor Daqus by PCR–DGGE. Letters in Applied Microbiology. 2011;53(2):134–40.

36. Jia L, Wei R, Jiang H, Yang XM, Xu YC, Shen QR. Application of bio-organic fertilizer significantly affected fungal diversity of soils. Soil Science Society of America Journal. 2010;74(6):2039.

37. Yu C, Hu X, Deng W, Li Y, Han G, Ye C. Soil fungal community comparison of different mulberry genotypes and the relationship with mulberry fruit sclerotiniosis. Sci Rep. 2016;6:28365.

38. Suleiman A, Lourenço KS, Pitombo LM, Mendes LW, Roesch L, Pijl A, et al. Recycling organic residues in agriculture impacts soil-borne microbial community structure, function and N2O emissions. Science of the Total Environment. 2018;631-632.

39. She S, Niu J, Chao Z, Xiao Y, Wu C, Dai L, et al. Significant relationship between soil bacterial community structure and incidence of bacterial wilt disease under continuous cropping system. Archives of Microbiology. 2017;199(2):1–9.

40. Mazzola M, Reynolds MP. Management of resident soil microbial community structure and function to suppress soilborne disease development. Climate Change & Crop Production. 2010;200–219.

41. Enwall K, Philippot L, Hallin S. Activity and composition of the denitrifying bacterial community respond differently to long-term fertilization. Acta Ecologica Sinica. 2009;71(12):8335–43.

42. Dignam BEA, O’Callaghan M, Condron LM, Kowalchuk GA, Nostrand JDV, Zhou J, et al. Effect of land use and soil organic matter quality on the structure and function of microbial communities in pastoral soils: Implications for disease suppression. PLoS One. 2018;13(5):e0196581.

43. Opina NL, Santiago SE, Tiongco RL, Miller S, Maghirang RG. Influence of cultural practices, host resistance and grafting on the incidence of bacterial wilt on eggplant. European Spine Journal. 2001;23(4):754–61.

44. Matsuda T. Fundamental studies on the breeding of bacterial wilt resistant varieties in tobacco. Bull Utsunomiya Tobacco Exp Stn. 1977;15:1–95.

45. Lit MTC, Opina NL, Lapiz RV, editors. Identification of eggplant varieties resistant to the leafhopper, shoot/fruit borer and bacterial wilt. Australian & New Zealand Journal of Psychiatry. 2002;45(12), 1061–8.

46. Gopalakrishnan TR, Singh PK, Sheela KB, Shankar MA, Kutty PCJ, Peter KV, et al. Development of bacterial wilt resistant varieties and basis of resistance in eggplant (Solanum melongena L.). Bacterial wilt disease and the Ralstonia solanacearum species complex. 2005;293–300.

47. Chae SY, Yang EY, Park EJ, Kim OR, Park TS, Sang GK, et al. Germplasm Evaluation and Selection for Breeding of Resistant Rootstock Varieties to Bacterial wilt in Pepper (Capsicum annuum L.). Science of horticulture volume. 2017;10: 114–115.

48. Irikiin Y, Nishiyama M, Otsuka S, Senoo K. Rhizobacterial community-level, sole carbon source utilization pattern affects the delay in the bacterial wilt of tomato grown in rhizobacterial community model system. Applied Soil Ecology. 2006;34(1):27–32.

49. Yamazaki H, Kikuchi S, Hoshina T, Kimura T. Effect of calcium concentration in nutrient solution on development of bacterial wilt and population of its pathogen Ralstonia solanacearum in grafted tomato seedlings. Soil Science & Plant Nutrition. 2000;46(2):535–9.

50. Hiromichi Yamazaki, Osamu Ishizuka, Tsuguo Hoshina. Relationship between resistance to bacterial wilt and nutrient uptake in tomato seedlings. Soil Science & Plant Nutrition. 1996;42(1):203–8.

51. Rui W, Zhang H, Sun L, Qi G, Shu C, Zhao X. Microbial community composition is related to soil biological and chemical properties and bacterial wilt outbreak. Scientific Reports. 2017;7(1):343.

